# Stochastic rank aggregation for the identification of functional neuromarkers

**DOI:** 10.1101/329383

**Authors:** Paola Galdi, Michele Fratello, Francesca Trojsi, Antonio Russo, Gioacchino Tedeschi, Roberto Tagliaferri, Fabrizio Esposito

## Abstract

**Background and aims:** The main challenge in analysing functional magnetic resonance imaging (fMRI) data from extended samples of subject (N>100) is to extract as much relevant information as possible from big amounts of noisy data. When studying neurodegenerative diseases with resting-state fMRI, one of the objectives is to determine regions with abnormal background activity with respect to a healthy brain and this is often attained with comparative statistical models applied to single voxels or brain parcels within one or several functional networks. In this work, we propose a novel approach based on clustering and stochastic rank aggregation to identify parcels that exhibit a coherent behaviour in groups of subjects affected by the same disorder and apply it to default-mode network independent component maps from resting-state fMRI data sets.

**Methods:** Brain voxels are partitioned into parcels through k-means clustering, then solutions are enhanced by means of consensus techniques. For each subject, clusters are ranked according to their median value and a stochastic rank aggregation method, *TopKLists*, is applied to combine the individual rankings within each class of subjects. For comparison, the same approach was tested on an anatomical parcellation.

**Results:** We found parcels for which the rankings were different among control subjects and subjects affected by Parkinson’s disease and amyotrophic lateral sclerosis and found evidence in literature for the relevance of top ranked regions in default-mode brain activity.

**Conclusions:** The proposed framework represents a valid method for the identification of functional neuromarkers from resting-state fMRI data, and it might therefore constitute a step forward in the development of fully automated data-driven techniques to support early diagnoses of neurodegenerative diseases.

## 1. Introduction

The recent advances in functional magnetic resonance imaging (fMRI) have made available high-quality data characterized by ever higher resolution images and shorter repetition times. This translated into an explosion of the dimensionality of data, thus generating the need of analysis techniques able to cope with the increased complexity of the problem.

To investigate how information is represented in the brain of healthy subjects, and how neurodegenerative diseases affect the physiological mechanisms underlying such a representation, a wide variety of statistical and computational methods have been applied to extract meaningful patterns of neural activity from fMRI data.

In population-level analysis, brain voxels (volumetric pixels) are usually analysed in isolation with traditional univariate techniques such as t-test or ANOVA. However, when the number of variables is huge, as is the case with fMRI data, issues such as the number of false positive and the multiple comparisons problem need to be addressed (see, e. g, Eklund et al. 2016). Moreover, voxels are features deriving from a geometrical representation of the brain that does not reflect the actual organization of neurons. While this might not constitute an issue in single-subject study, in a cohort study inter-subject variability hinders the generalizability of the results: in fact, a voxel found to be significant on a given subject may not be significant on a different subject, or even fall in a different brain region.

When conducting population studies, the goal is to analyse group-specific behaviour starting from the product of a first-level analysis consisting in single-subject activation maps. One of the limitations of this approach is that, due to the low signal-to-noise ratio of single-subject images, and to within and between subject variability, the measurable effects might be small or masked by noise. For this reason, in this work we propose an approach based on similarity within groups as opposed to traditional comparative approaches that focus on searching discriminative patterns.

We present a new framework based on clustering and stochastic rank aggregation methods to identify functional neuromarkers starting from single-subject activation maps. Clustering has been previously applied in fMRI data analysis to extract patterns from raw time series (Goutte 1999) or from second level features extracted from data (Goutte et al. 2001), and in group level analyses (Thirion et al. 2006; Heuvel et al. 2008). In combination with single-subject independent component analysis (ICA) (Hyvärinen and Oja 2000), clustering has been also used to identify the most similar ICA components within a single group of subjects (Esposito et al. 2005), but to the best of our knowledge this is the first time that it is used in combination with rank aggregation to identify regions of the brains that show a common behaviour among subjects belonging to the same class or group and investigate the differences between different classes or groups. Since this approach is fully data-driven, there is no need for strong assumptions about data distribution as in parametric models, and can therefore be equally applied to, e. g., ICA maps derived from resting-state fMRI data or conventional activation maps derived from a general linear model analysis of activation time-courses. Nor it is necessary to specify in advance regions of interest, since a small subset of informative regions automatically emerges from the analysis. This framework was tested on a real-world data set consisting of individual ICA-derived default-mode network (DMN) maps from resting-state fMRI scans of healthy controls and of subjects affected by amyotrophic lateral sclerosis (ALS) and Parkinson’s disease (PD). Although very different in their clinical manifestations, both these pathologies affect the motor system. However, previous structural and functional studies have reported widespread extra-motor changes in patients with ALS, with frontal deficits linked to neuropsychological impairment and differences in the activation of the default mode and sensorimotor network (Lomen-Hoerth et al. 2003; Kiernan et al. 2011; Luo et al. 2012; Verstraete et al. 2014; Chiò et al. 2014; Verstraete and Foerster 2015; Menke et al. 2017). Network-based studies on PD revealed altered connectivity in the sensorimotor network, DMN, fronto-parietal network and temporal-occipital networks (Gattellaro et al. 2009; Olde Dubbelink et al. 2014; Tessitore et al. 2014; Suo et al. 2017; Herrington et al. 2017; Hepp et al. 2017). The choice of validating the proposed framework using the DMN is aimed at facilitating the interpretation of the results. The DMN is by far the most studied resting-state network, it appears to be the most stable across subjects both in health and in disease, it is known to interact with higher order networks and can be considered a cognitive baseline for a subject (Greicius et al. 2003; Damoiseaux et al. 2006; Harrison et al. 2008; Buckner et al. 2008; Tedeschi and Esposito 2009; Whitfield-Gabrieli and Ford 2012; Raichle 2015). Additionally, previous works have investigated the role on the DMN in association with ALS (Mohammadi et al. 2009; Tedeschi et al. 2012; Agosta et al. 2013; Iyer et al. 2015; Trojsi et al. 2015) and PD (Van Eimeren et al. 2009; Tessitore et al. 2012b; Gorges et al. 2013; Amboni et al. 2015; Luo et al. 2015). On the other hand, higher order networks imply other clinical considerations related to the diseases under study. For example, the sensory-motor network is known to be suppressed in subjects affected by ALS and is characterized in general by a lower signal-to-noise ratio (Mohammadi et al. 2009; Tedeschi and Esposito 2009; Tedeschi et al. 2012) while in PD patients it has been shown that levodopa enhances the sensorimotor network functional connectivity in the supplementary motor area, a region where drug-naïve PD patients exhibit reduced signal fluctuations compared with untreated patients (Esposito et al. 2013). While the proposed method can be applied to any network, a study on sensory/motor function should consider also other clinical factors that can potentially affect the results and consequently their interpretation.

## 2. Methods

### 2.1. Ethics statement

The institutional review board for human subject research at the Università degli Studi della Campania “Luigi Vanvitelli” approved the study and all subjects gave written informed consent before the start of the experiments.

### 2.2. Participants

Data come from a cohort of 121 subjects, with age ranging from *38* to *82* years (mean age *63.87 ± 8.2*). Specifically, they include *41* patients (*20* women) from a clinical study on amyotrophic lateral sclerosis (Tedeschi et al. 2012); *37* patients (*14* women) from a clinical study on Parkinson’s Disease (Tessitore et al. 2012a; Tessitore et al. 2012b; De Micco et al. 2013; Amboni et al. 2015) and *43* control subjects (*23* women) from the same clinical studies. All control subjects were free from (and had no history of) neurological disorders.

### 2.3. MRI Data Acquisition and Pre-processing

A 3T scanner equipped with an 8-channel parallel head coil (General Electric Healthcare, Milwaukee, Wisconsin) was used for the acquisition of MRI images.

A sequence of 240 volume was acquired using gradient-echo T2* -weighted MR imaging (TR = 1508ms, axial slices = 29, matrix = 64 × 64, field of view = 256mm, thickness = 4mm, inter-slice gap = 0mm).

Subjects were asked to stay awake and motionless and to keep their eyes closed during the scans.

To register and normalize fMRI images, high resolution T1-weighted sagittal images were acquired in the same session (GE sequence IR-FSPGR, TR = 6988ms, TI = 1100ms, TE = 3.9ms, flip angle = 10, voxel size = 1mm × 1mm × 1.2mm).

Data pre-processing was performed with BrainVoyager QX (Brain Innovation BV, Maastricht, the Netherlands) including slice timing correction, 3D rigid body motion correction and high-pass temporal filtering. After registration to structural images, functional images were normalized to fit the Talairach standard space using a 12-parameter affine transformation and resampled to an isometric 3mm grid covering the entire Talairach box. Finally, all volumes were visually inspected to assess the impact of geometric distortion on the final images, which was judged to be negligible for a whole-brain analysis.

For each subject, 40 independent components (ICs) were extracted with the fastICA algorithm (Hyvarinen 1999), accounting for more than 99.9% of the total variance. The number of ICs corresponds to one sixth of the number of time points (Greicius et al. 2007; Bosch et al. 2018). Among the extracted components, the one associated with the DMN was selected as the one with the highest goodness of fit (GoF) with a DMN mask^1^ from a previous study on the same MRI scanner with the same protocol and pre-processing (Esposito et al. 2010), where the GoF is defined as the difference between the average component score inside the mask minus the average component score outside the mask (Greicius et al. 2004; Esposito and Goebel 2011). To avoid ICA sign ambiguity, each component sign was adjusted to have all GoF values as positive (see, e. g., (Esposito et al. 2008)).

The DMN mask used for this study was originally obtained from a self-organizing group-level ICA (sogICA) analysis (Esposito et al., 2005) on a different group of subjects by applying a voxel-level threshold of p=0.05 (Bonferroni corrected for multiple voxel-level comparisons). Thus, following (Smith et al. 2009), 2009, to select (and validate) the DMN mask, the same threshold was applied to all original sogICA components and the resulting masks were applied to an externally validated (“well-matched”) DMN component templated available on line (https://www.fmrib.ox.ac.uk/datasets/brainmap+rsns/). The best matching between the sogICA component mask and the external well-matched DMN component map resulted from the highest average ICA score of the latter among all candidate masks.

### 2.4. Overview of the methodology

Following the pre-processing and the extraction of the DMN from each subject’s data (as described in section 2.3), the input data consist of a matrix *N* ×*V* where *N* is the number of subject and *V* is the number of voxels, and each entry (*i, j*) represents the contribution of voxel *j* to the DMN map of subject *i*.

First, voxels are partitioned into parcels using one of the two methodologies detailed in the following section. The parcellation is applied to the data of each subject and a representative feature (the median) is selected for each parcel. A ranking is then computed for each subject by sorting in descending order the extracted features. Finally, rankings are aggregated by class of subjects through stochastic rank aggregation (section 2.6); the goal is obtaining a subset of brain regions that share a common behaviour within classes. See Figure 1 for a graphic overview of the methodology.

**Figure 1.**
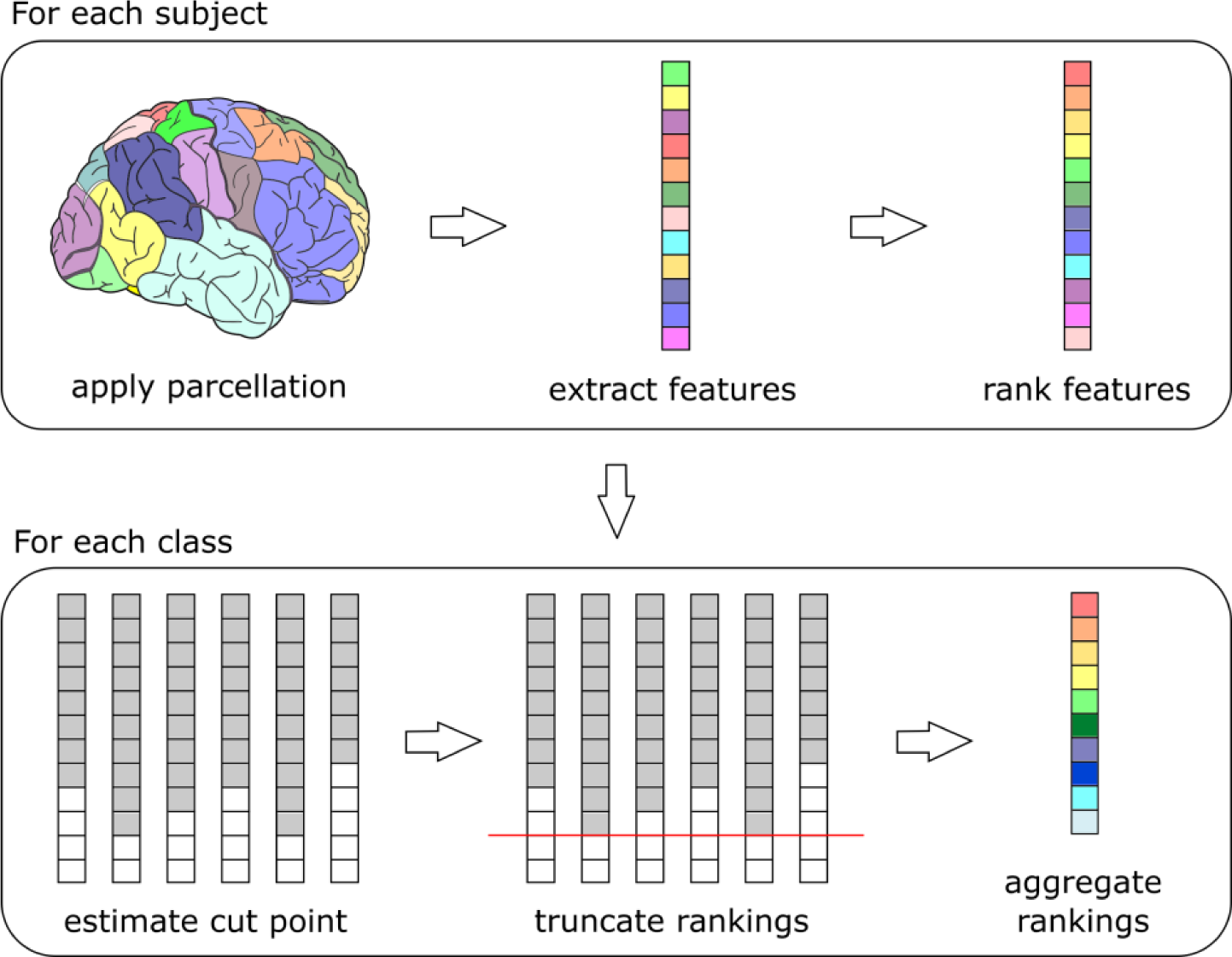
Overview of the methodology. A parcellation (derived by a clustering algorithm or from an atlas) is applied to each subject’s whole-brain ICA map. From each parcel, a representative feature is extracted (e.g., the median value). Features are ranked in descending order. In this way, each subject expresses a ranking of the parcels. Then, a stochastic rank aggregation method (TopKLists) is applied to the ensemble of rankings of each class of subjects. This method first estimates the rank position beyond which concordance among rankings degenerates into noise (cut point), then truncates all rankings at this position, and finally computes an aggregate ranking using a Cross-entropy Monte Carlo method.

### 2.5. Brain parcellation

Working on brain parcels instead of single voxels is convenient for many reasons. As mentioned above, brain activity is likely to span over multiple voxels. Therefore, the aggregation of several voxels in a single agglomerated feature may reduce redundancy and improve signal-to-noise ratio, and this could in turn increase the prediction accuracies of learning models.

To validate our method for the selection of relevant regions of the brain based on stochastic rank aggregation, we adopted two different approaches to brain parcellation: the former based on anatomical information and the latter based on a data-driven clustering technique.

Anatomical parcellations are derived from an atlas that defines brain regions on a template image. The one used in this work is the Automated Anatomical Labelling (AAL) atlas (Tzourio-Mazoyer et al. 2002) that consists of 90 parcels. Since this approach is data-independent, it allows an objective comparison of the results of different models.

Clustering methods can be applied after the standard pre-processing of fMRI data to segment data in groups maximizing a given measure of intra-group similarity. We adopted k-means clustering and Pearson’s correlation coefficient as measure of similarity between voxels across subjects. Moreover, to obtain more stable and reliable sets of features, clustering solutions were enhanced through consensus techniques, i.e. different clustering solutions were combined into a final clustering to improve the quality of individual data clusterings, as described in a previous work (Galdi et al. 2017). The consensus-based approach can be briefly described as follows: 1) first, several partitions of voxels are generated with k-means with random initialization, to ensure variability among base solutions; 2) a voxel-similarity matrix (consensus matrix) is built by counting how many times each pair of voxels is assigned to the same cluster across partitions; 3) a consensus clustering solution is obtained by applying a hierarchical clustering algorithm that uses the consensus matrix as a similarity matrix; 4) finally, the information contained in the consensus matrix is used to perform a further feature selection step, by retaining only a stable subset of voxels that are consistently assigned to the same groups. This is attained by applying two thresholds: σ, that defines how often two features should be clustered together to be considered a stable pair, and ν, that is the minimum number of stable pairs a voxel needs to be part of to be selected. These two thresholds are tuned on the data. In our previous work, the number of clusters (k=500) was chosen heuristically to guarantee that clusters were sufficiently big to be easily mapped to an anatomical region for a posteriori validation. After the application of the consensus-based voxel filtering, the final solution consists of 405 brain parcels. Since part of the data were used as a training set to obtain the consensus solution, the final clustering was applied on a test set to extract the features for the subsequent steps of the analysis.

To demonstrate that our consensus-based approach identified a stable solution, we generated 100 bootstrap samples of the original dataset (sampling with replacement 2/3 of the initial set of subjects) and repeated the whole procedure on each sample (von Luxburg 2010), varying the number of clusters k ∈{90, 100, 200, 300, 400, 500}. We then computed the adjusted normalized mutual information (NMI) between each pair of partitions to assess the stability of the resulting clustering solutions (Meilă 2007; Vinh et al. 2009; Vinh and Epps 2009). Here, we have chosen the adjusted version of the NMI score because it corrects the effect of agreement solely due to chance and it accounts for the fact that the NMI is generally higher for two clusterings with a larger number of clusters, regardless of whether there is actually more information shared (Vinh et al. 2009; Amelio and Pizzuti 2015).

Results are shown in figure 2, before and after the consensus-based voxel filtering. We can observe that the NMI score increases as the number of cluster increases, and the application of the consensus-based filter improves the score in every test. Note that after the application of the filter, some voxels are discarded from each partition, hence the NMI score is computed only on the intersection of voxels between each pair of partitions. Figure 3 shows the percentage of voxels that (on average) are selected by the filter (in yellow), and the (average) percentage of filtered voxels that are in the intersection between two partitions. This latter value is an indirect measure of the stability of the filtered solutions. Increasing the number of clusters in the partitions leads to a decreasing percentage of filtered voxels, indicating that the base solutions were indeed less stable. However, the consensus solution that we adopted (k=500, post-filtering) is the one with the highest NMI score, and the average percentage of voxels in the intersection is still reasonably high (∼60%).

**Figure 2.**
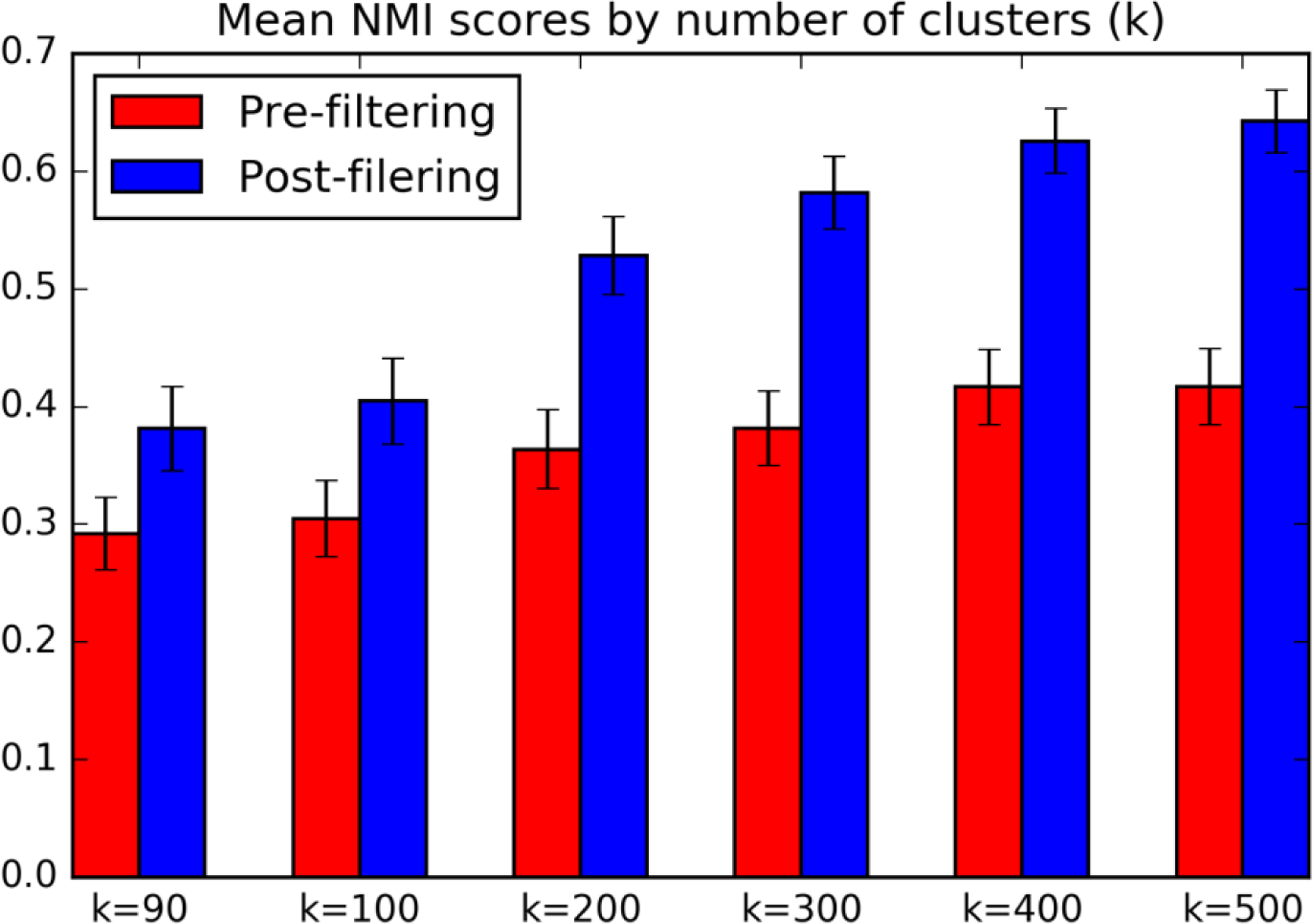
Results of clustering stability assessment for different number of clusters (k). The adjusted NMI score was computed between pairs of partitions obtained on 100 bootstrap samples. The bars represent the mean adjusted NMI score, before the application of the consensus-based filter (in red) and after (in blue).

**Figure 3.**
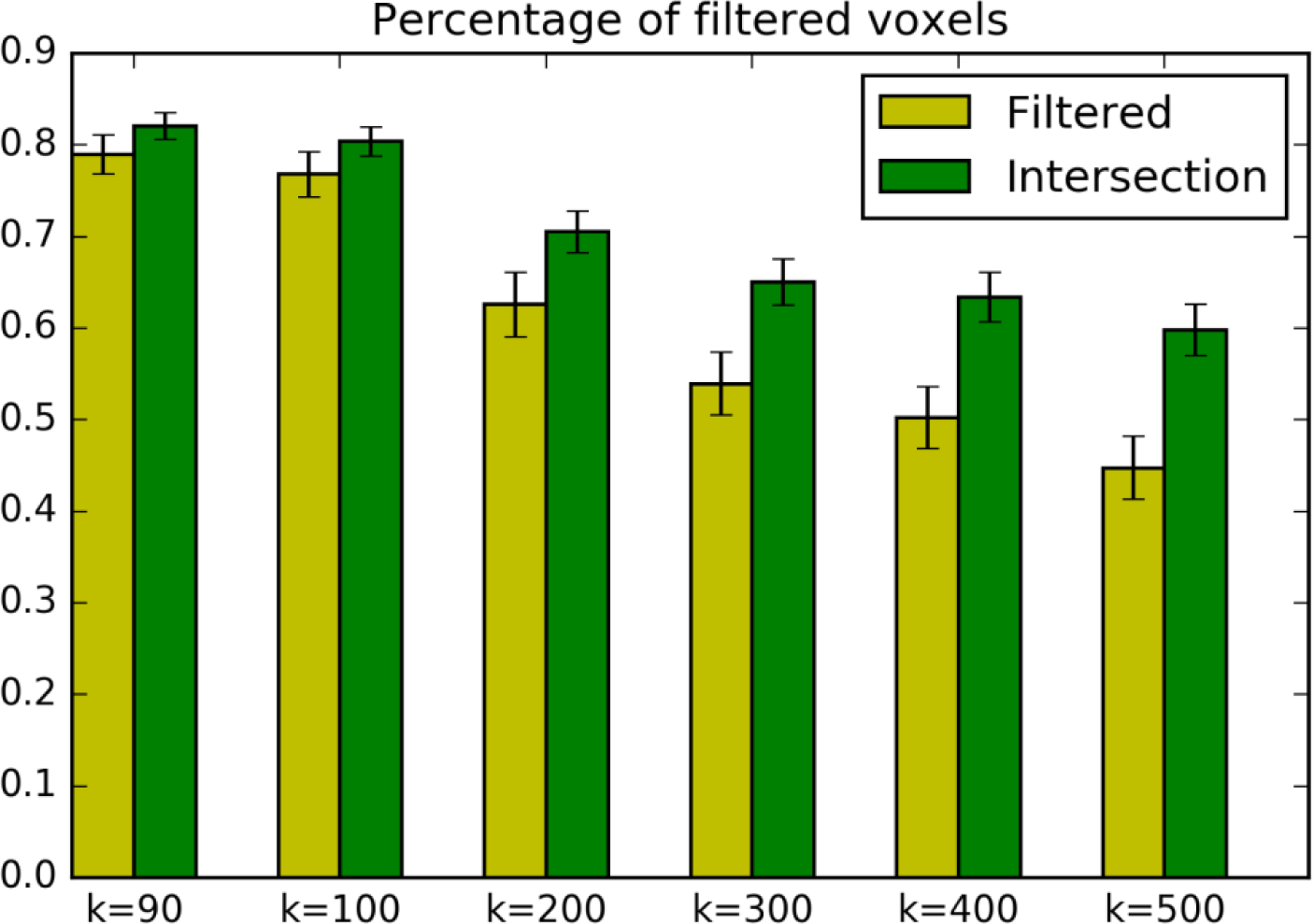
Average percentage of voxels selected by the consensus-based filter across bootstrap samples (in yellow). Average percentage of shared voxels after the application of the consensus-based filter between pairs of partitions obtained on 100 bootstrap samples (in green).

Once voxels are segmented in parcels, a representative feature can be extracted from each group to obtain a compressed representation of the input data. On the one hand this can be considered as a dimensionality reduction step, aimed to decrease the number of variables and thus facilitating the application of more sophisticated statistical models. On the other hand, aggregating the information described by several voxels allows working at a higher level of abstraction, that of brain regions, and this has multiple advantages: a) it captures the modular organization of the brain; b) it aids the generalization of the results since clusters of voxels are built across subjects; c) features are easier to interpret because they can be reliably mapped to brain regions. Compared to other dimensionality reduction techniques (e.g. PCA), clustering produces features that can be put into bijective correspondence with voxels, retaining information useful for visualization and interpretation of the discriminative features.

The main drawback of atlas-based approaches is that they are prone to errors in the segmentation of functional regions and this might translate into a decreased sensitivity of the models. Data-driven approaches are, by contrast, more sensitive to noise and since they are not based on a priori defined anatomical regions they require a further step to map features onto an atlas to allow a biological interpretation.

### 2.6. TopKLists

*TopKLists* (Schimek et al. 2012; Schimek et al. 2015) is a stochastic rank aggregation method that, starting from an ensemble of rankings of a set of items, outputs a new ranking of a subset of the same objects. It works by estimating the rank position *k* beyond which the concordance among the input rankings degenerates into noise. Once *k* has been computed, all rankings are cut at position *k,* thus selecting the top *k* elements of each ordered list (hence the name). Finally, all the sublists of length *k* are aggregated through a cross-entropy Monte Carlo method (Lin and Ding 2009).

The estimation procedure for the cut point can be summarized as follows. First, a cut point is estimated for each pair of ordered lists. For each pair, an indicator vector *I* is built, where *I*_*i*_= 1 if the item ranked *i* in the first list is ranked no more than *δ* positions away from rank *i* in the second list, and *I*_*i*_ = 0 otherwise. Namely, a value of 1 means that a given object was assigned a similar ranking in both lists. The indicator vectors *I*_1_, …, *I*_*N*_ are assumed to be independent Bernoulli random variables, with 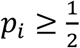 for *i* < *i*_0_ and 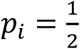 for *i* ≥ *i*_0_, where *p*_*i*_ is the probability that *I*_*i*_= 1 and *i*_0_ is the rank position where the consensus of the two lists breaks down and noise takes over^2^. In other words, before position *i*_0_, the probability that *I*_*i*_ = 1 is above chance level; after position *i*_0_, it is equal to chance level. For each vector *I*_*i*_, an estimate of *i*_0_ is computed with an iterative procedure, alternating a sequence of two steps. At each iteration, the algorithm updates the current estimate of the position *i*_0_, with the initial estimate being the first position of *I*_*i*_. Odd iterations start *rv* positions to the left of the current estimate, while even iterations start *rv* positions to the right of the current estimate. At iteration *s*_*j*_, a sample of size *v* is extracted consisting of elements *I*_*i*_ with *i* comprised among the first *v* indices to the right of 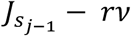, if *j* is odd, or to the left of 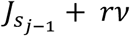, if *j* is even (where 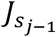 is the position where the previous iteration ended). Informally, at odd (even) iterations, the sample is extracted taking *v* consecutive items starting *rv* positions to the right (left) of the current estimated position (see Figure 4 for a schematisation of these stages). The sample mean is used to estimate the consensus probability *p_i_*: for even iterations 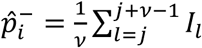 and for odd iterations 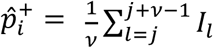. In even iterations, we move to the left by unitary steps until we reach the first point where 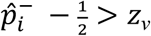. In odd iterations, we move to the right as long as the inequality 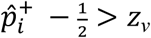 holds. The threshold *z*_*v*_ is defined as

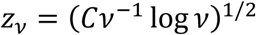

with 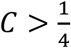, to control for moderate deviations in the degree of correspondence (Hall and Schimek 2012). The algorithm terminates when one of the following stop conditions is met: a) the algorithm enters a loop between two adjacent stages; b) for some *j*, 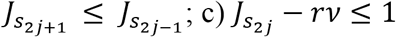. Once an estimate of *i*_0_ is computed for each pair of ordered lists, the final cut point is computed as the maximum of the estimated values.

**Figure 4.**
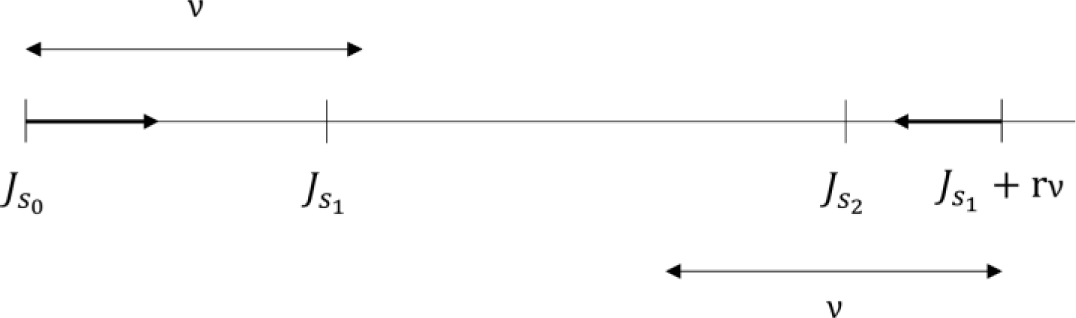
A schematisation of the *TopKList* algorithm. The first iteration starts at position 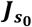 (the first entry of the indicator vector) and stops in position 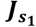, that is the first position where 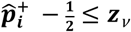. The second iteration starts at position 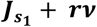 and ends in position 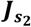, the first position where 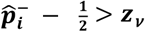.

In this work, *TopKLists* was applied to combine the rankings of brain parcels expressed by each subject of a class. Specifically, for each subject, the medians of the brain regions were computed and sorted in descending order, then all rankings of a class were aggregated in a single list, containing a subset of regions that were ranked similarly across subjects of the same class. This means that the concordance among rankings and subsequently the length of the final aggregate ranking are estimated using only data from subjects of the same class, hence for each class the number of retained regions can vary.

## 3. Results

Tables 1 and 2 report the regions selected by *TopKLists* for each class with the relative rankings for anatomical parcels and clusters, respectively (the number of voxels and the Talairach coordinates of the geometric mean is reported for each region).

**Table 1.** Ranking per class (AAL). The first two columns report the name and the number of the AAL ROI, for reference. Columns CTRL(30), ALS(27) and PD(26) report the ranking for each parcel for controls, ALS and PD patients, respectively. The number between brackets indicates the number of top ranking regions selected per each class. The last two columns report the number of voxels and the coordinates of the geometric mean (in Talairach space) for each parcel.

| AAL ROI | AAL # | CTRL(30) | ALS(27) | PD(26) | # voxels | geom. mean |
| --- | --- | --- | --- | --- | --- | --- |
| Frontal_Sup_L | 3 | 15 | 19 | 17 | 1402 | (14.4,12.2,29.9) |
| Frontal_Sup_R | 4 | 22 | 22 | 23 | 1513 | (15.8,11.7,17) |
| Frontal_Mid_L | 7 | 27 | 25 | 22 | 1765 | (15.5,14,34.7) |
| Frontal_Mid_R | 8 | 24 | - | - | 1866 | (15.6,14.4,11.8) |
| Frontal_Sup_Medial_L | 23 | 6 | 3 | 6 | 1099 | (10.1,14.8,25.5) |
| Frontal_Sup_Medial_R | 24 | 3 | 9 | 13 | 853 | (9.7,14.8,21) |
| Frontal_Med_Orb_L | 25 | 1 | 2 | 2 | 300 | (7.8,26.5,25.7) |
| Frontal_Med_Orb_R | 26 | 4 | 11 | 11 | 367 | (8.9,26.4,21.4) |
| Rectus_L | 27 | 25 | - | - | 374 | (13.2,30.2,25.8) |
| Rectus_R | 28 | 28 | - | - | 352 | (13.9,30.2,21.3) |
| Cingulum_Ant_L | 31 | 16 | 17 | 14 | 590 | (14.6,20.3,25.4) |
| Cingulum_Ant_R | 32 | 11 | 18 | 21 | 537 | (14.1,19.8,21.3) |
| Cingulum_Mid_L | 33 | - | - | 26 | 674 | (30.9,13.2,26.1) |
| Cingulum_Post_L | 35 | 2 | 1 | 1 | 205 | (38.7,19.1,26.1) |
| Cingulum_Post_R | 36 | 5 | 4 | 5 | 152 | (38.4,20,22.1) |
| ParaHippocampal_L | 39 | 26 | - | - | 410 | (30,32.3,31.2) |
| ParaHippocampal_R | 40 | 29 | - | - | 471 | (29.8,32.1,16.2) |
| Calcarine_L | 43 | 7 | 8 | 10 | 866 | (50.1,25.6,27) |
| Calcarine_R | 44 | 8 | 10 | 15 | 743 | (48.8,24.5,19.5) |
| Cuneus_L | 45 | 9 | 6 | 7 | 579 | (51,19.3,26.7) |
| Cuneus_R | 46 | 20 | 16 | 18 | 577 | (51.1,19,20.2) |
| Lingual_L | 47 | 19 | 23 | - | 868 | (46.4,28.9,29.3) |
| Lingual_R | 48 | 21 | 24 | - | 940 | (46.2,28.5,19.3) |
| Occipital_Sup_L | 49 | - | 27 | 25 | 554 | (52.5,19.1,29.9) |
| Occipital_Sup_R | 50 | - | 26 | 24 | 573 | (51.7,18.2,16.8) |
| Occipital_Mid_L | 51 | 12 | 15 | 12 | 1222 | (50.8,22.8,34.9) |
| Occipital_Mid_R | 52 | 18 | 14 | 16 | 789 | (51.1,21.4,12.5) |
| Angular_L | 65 | 23 | 12 | 4 | 461 | (44.7,16.3,38.5) |
| Angular_R | 66 | 30 | 20 | 9 | 679 | (45.1,15.1,9.8) |
| Precuneus_L | 67 | 10 | 5 | 3 | 1308 | (43.5,12.5,26.9) |
| Precuneus_R | 68 | 13 | 7 | 8 | 1257 | (43.5,13.8,21.2) |
| Temporal_Mid_L | 85 | 17 | 21 | 20 | 1604 | (35.6,27.1,41.8) |
| Temporal_Mid_R | 86 | 14 | 13 | 19 | 1542 | (37.3,26.7,5.9) |

**Table 2.** Ranking per class (clusters). The first column reports numeric identifiers of clusters, for reference. Columns CTRL(8), ALS(9) and PD(8) report the ranking for each cluster for controls, ALS and PD patients, respectively. The number between brackets indicates the number of top ranking clusters selected per each class. The fourth column indicates in which Brodmann area each cluster fall. The column AAL AREA indicates the corresponding parcel in the anatomical based solution; when more than one region is reported, the clusters lie on the boundary of parcels. The last two columns report the number of voxels and the coordinates of the geometric mean (in Talairach space) for each parcel.

| Cluster | CTRL(8) | ALS(9) | PD(8) | Brodmann Area | AAL Area | # voxels | geom. mean |
| --- | --- | --- | --- | --- | --- | --- | --- |
| 86 | - | - | 5 | 40, 41 | - | 27 | (31,22,4) |
| 95 | - | - | 6 | 40 | - | 51 | (37,14.6,7.1) |
| 98 | 2 | 3 | 3 | 10 | 25, 27 | 81 | (8,28,27) |
| 160 | - | - | 8 | 40 | - | 71 | (40,12.3,9.1) |
| 182 | 3 | 2 | 4 | Anterior cingulate (WM) | - | 27 | (13,25,28) |
| 197 | 4 | - | 1 | 22 (WM) | - | 61 | (38.6,18.4,4.2) |
| 223 | - | 7 | - | 23, 24 | 31 | 81 | (25,16,25) |
| 265 | 1 | 1 | 2 | 10, 32 | 25 | 54 | (8.5,25,25) |
| 297 | - | - | 7 | 39 | 65, 85 | 54 | (41.5,20.5,37) |
| 314 | - | 9 | - | 7, 18 | 67,45 | 54 | (47.5,16,25) |
| 341 | 7 | 8 | - | WM | - | 27 | (34,19,31) |
| 380 | 6 | - | - | 21 | 86 | 101 | (31.2,28,5.7) |
| 381 | - | 5 | - | 22 | - | 17 | (28.1,24.9,4.5) |
| 382 | 8 | - | - | 20, 37 | 86 | 98 | (37,31.4,4) |
| 383 | - | 6 | - | 37 | 86 | 72 | (40.8,28,5.3) |
| 386 | 5 | - | - | 22, 47 | - | 78 | (18,26.9,8.1) |
| 425 | - | 4 | - | 8 | 3, 7 | 26 | (13,13,31) |

The number of anatomical areas selected per class (30 for controls, 27 for ALS, 26 for PD) was higher than the number of selected clusters (8 for controls, 9 for ALS, 8 for PD) albeit the size of functional clusters was smaller than the size of anatomical areas. Figure 5 represents with Venn’s diagrams the overlap of regions and clusters shared among classes. There are more anatomical areas shared among all three classes than class-specific anatomical areas; conversely, there are more class-specific than shared clusters. For the ALS group, there are 5 class-specific parcels in the clustering-based ranking while there are none in the anatomical-based one.

**Figure 5.**
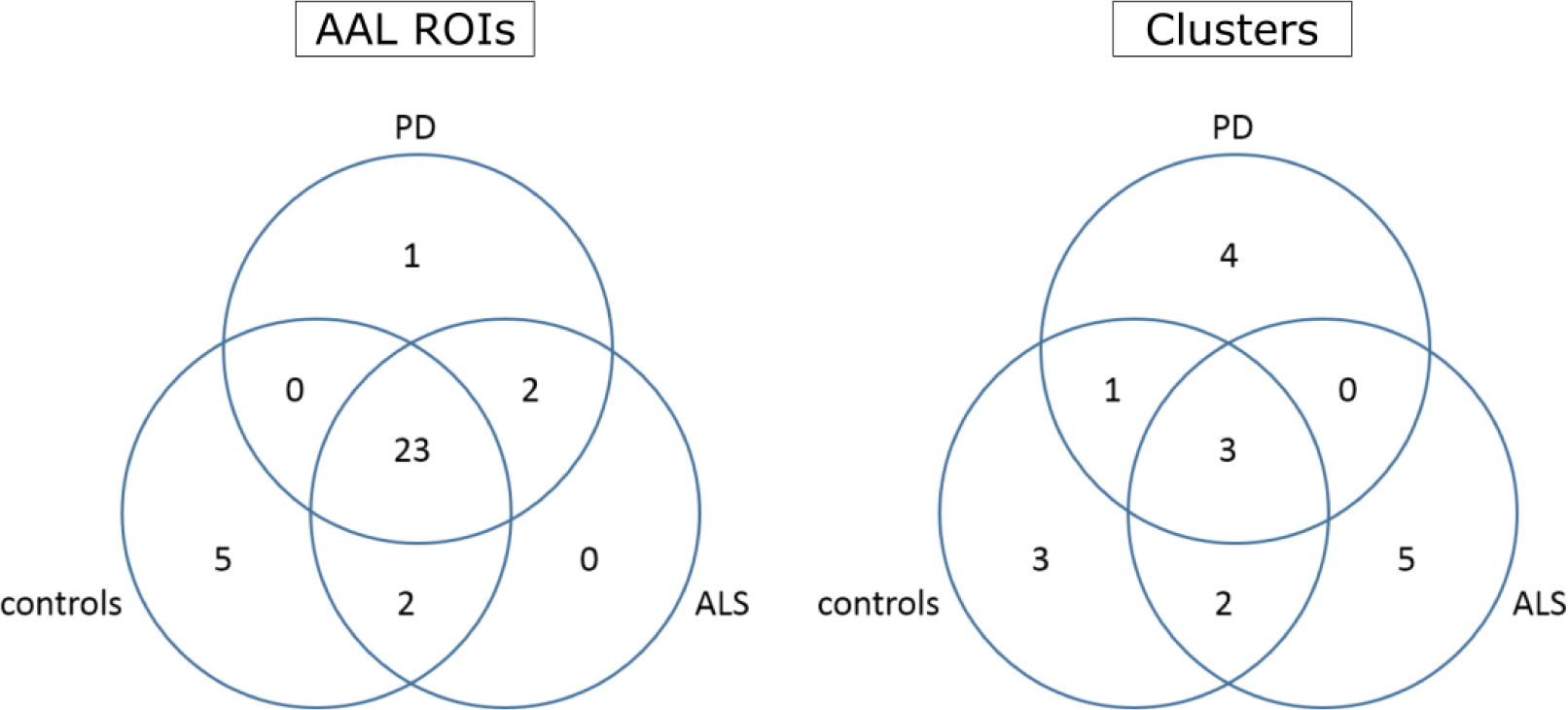
Venn diagram of the selected anatomical areas (left) and of selected clusters (right). Most of the anatomical areas are shared among the three classes, while there are more class-specific regions when using clusters. For the ALS group, there are 5 class-specific parcels in the clustering-based ranking while there are none in the anatomical based one.

Figures 6 and 7 show a comparison between the anatomical and the clustering-based solutions in corresponding regions, on an MRI image and on an inflated representation of the cortex. Cluster 265 and AAL area 25 (corresponding to the left medial orbitofrontal cortex) are both in the first or the second rank in all three classes (figure 6). Both parcels identify approximately the same region, but the functional cluster is smaller and has a substantial overlap with Brodmann area 10.

**Figure 6.**
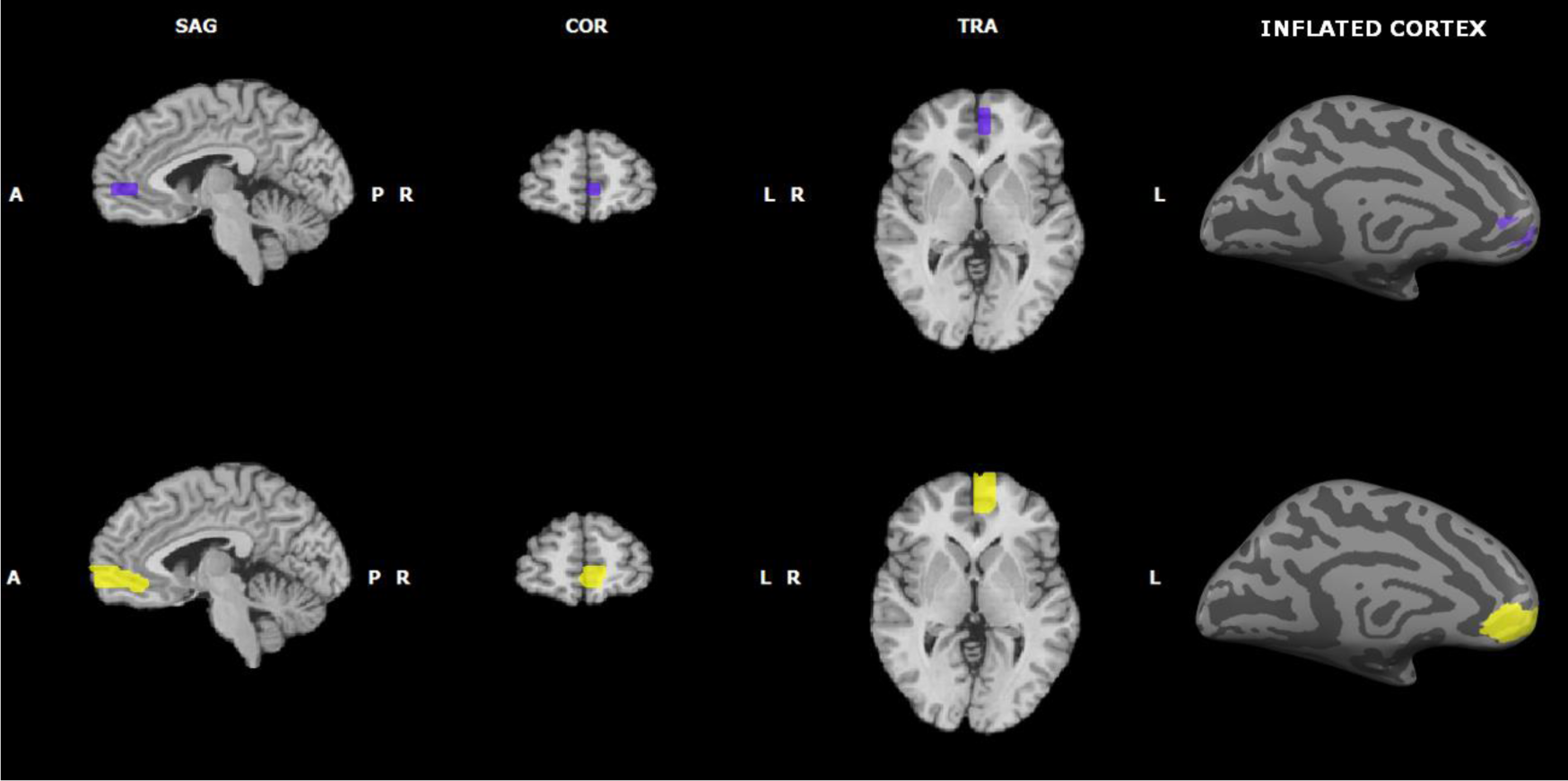
Comparison between anatomical areas and clusters in corresponding regions superimposed on a standard MRI image and on an inflated cortex. The top row shows cluster 265 while the bottom row shows AAL ROI 25

**Figure 7.**
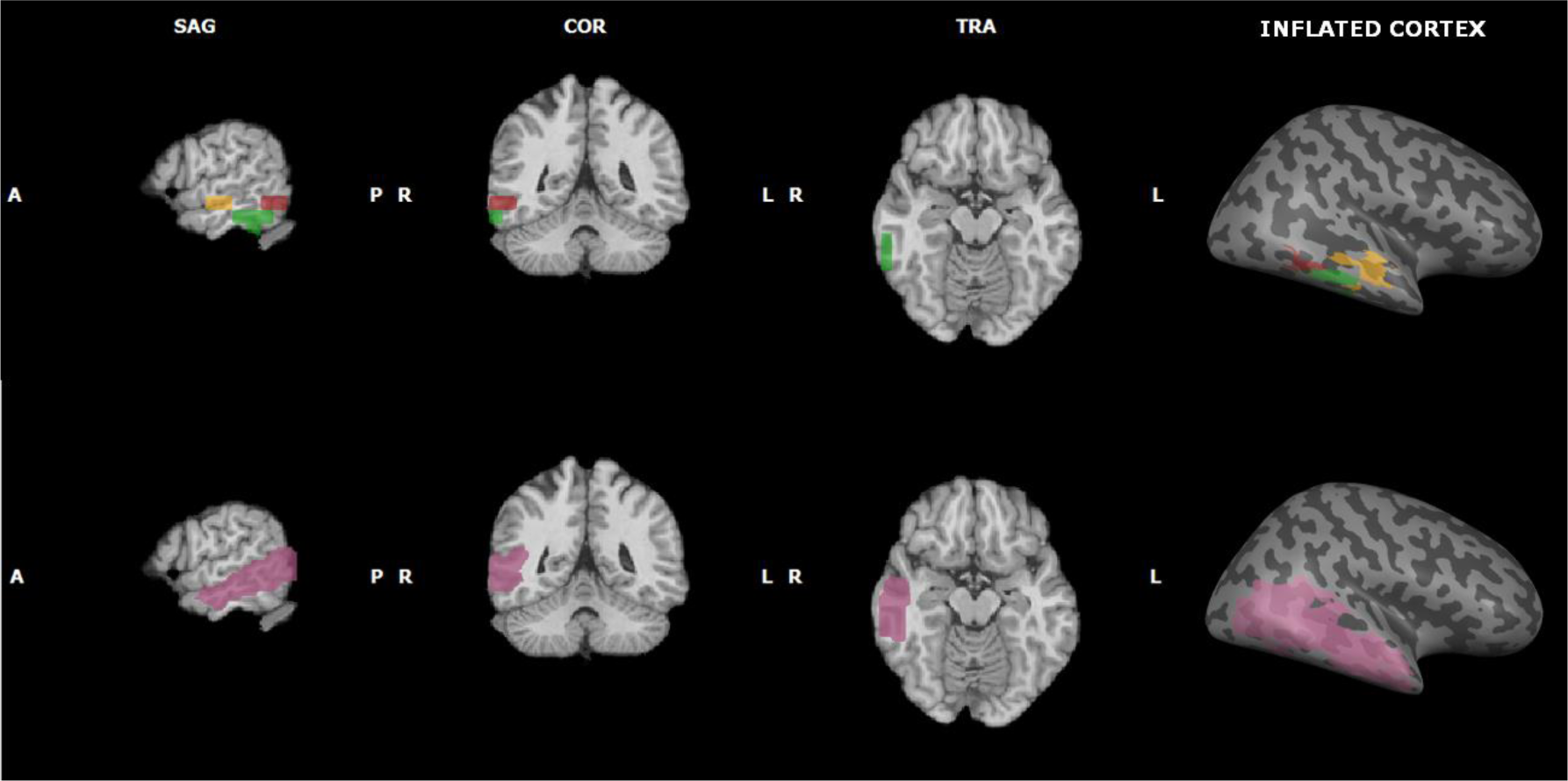
Comparison between anatomical areas and clusters in corresponding regions superimposed on a standard MRI image and on an inflated cortex. The top row shows clusters 380 (in orange), 382 (in green) and 383 (in red). The bottom row shows AAL ROI 86.

Based on their respective rankings, AAL region 86 (right middle temporal gyrus) can be compared with functional clusters 380, 382 and 383 (figure 7). In this case, the AAL parcel is present in the top rankings of all classes, while clusters 380 and 382 are specific for the control class, and cluster 382 is specific to the ALS class.

Figures 8 to 11 represent with a box-plot the distribution of the median of each of the selected regions across subjects of the same class (for anatomical regions in figures 8 to 10 and for clusters in figure 11), with the parcels ordered according to the rankings. In most cases, class specific regions (black box-plots) exhibit less variability than others,. In the anatomical-based solution for the control class (figure 8), the five class-specific regions occupy the highest ranks and exhibit narrow distributions; all of the remaining regions are shared among all classes except for two that are shared only with the ALS class and are present in the first half of the ranking. The only class-specific region for PD occupies the first rank, followed by the two regions shared with the ALS class (figure 9). The remaing regions are shared with all classes and show a high variability across subjects (figure 9). For what concerns the anatomical areas relative to the ALS class (figure 10), there are no class-specific regions, but the ones shared with PD and controls, respectively, are listed in the top ranks with narrow distributions. Also in this case the regions shared among all classes exhibit a high variability. Considering now the clustering-based solutions (figure 11), we can see that in general the parcels show a reduced variability compared to the anatomical regions, but this is not unexpected since the clusters are smaller and the selected parcels are fewer. Another difference with the anatomical approach is that class-specific parcels occupy the second half of the ranking while the clusters shared by all classes (265, 98 and 182) are in the top ranks.

**Figure 8.**
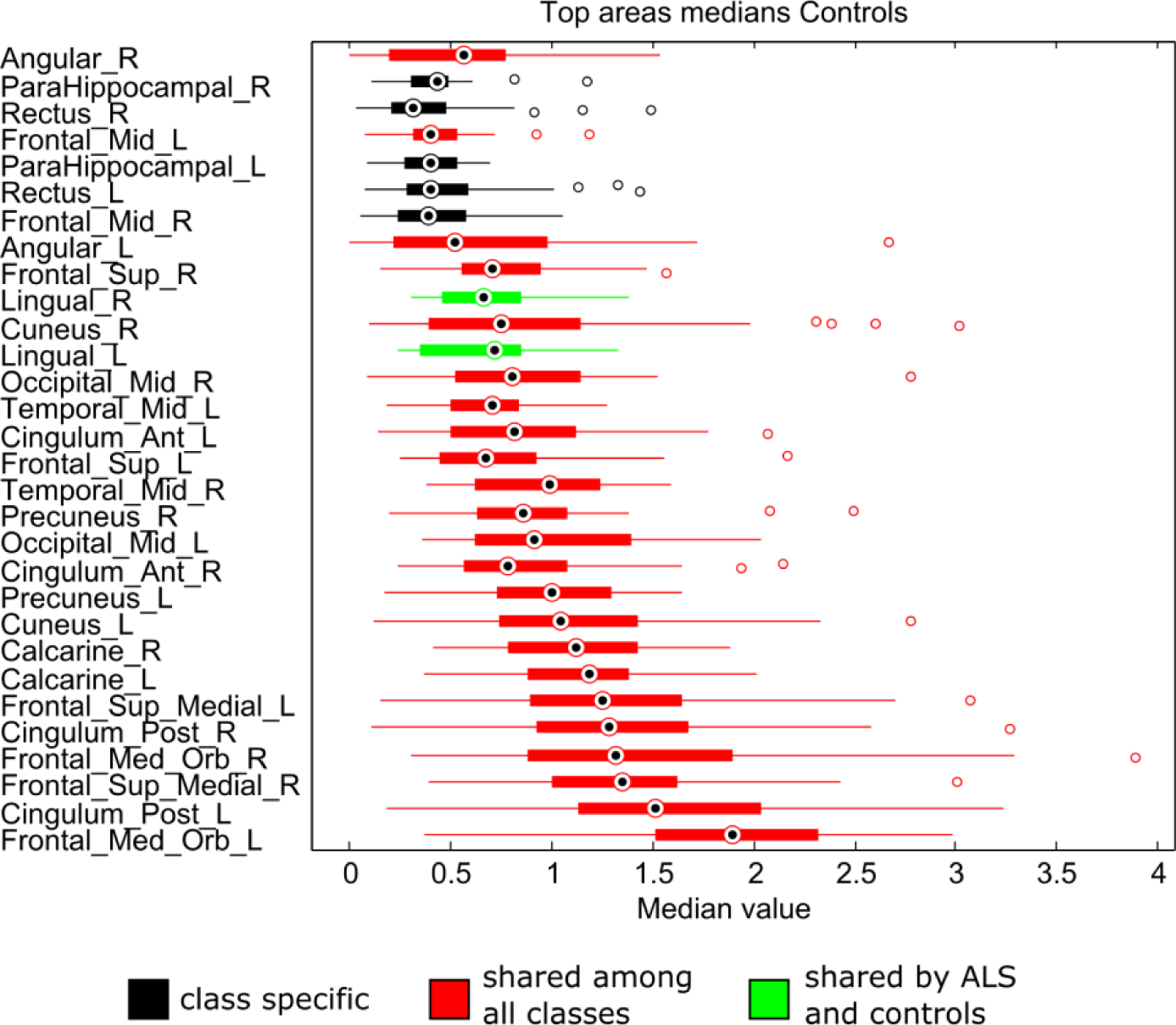
Box plots of the distribution of the medians of each of the selected anatomical areas across subjects of the control class. The five class-specific regions (in black) occupy the highest ranks and exhibit narrow distributions.

**Figure 9.**
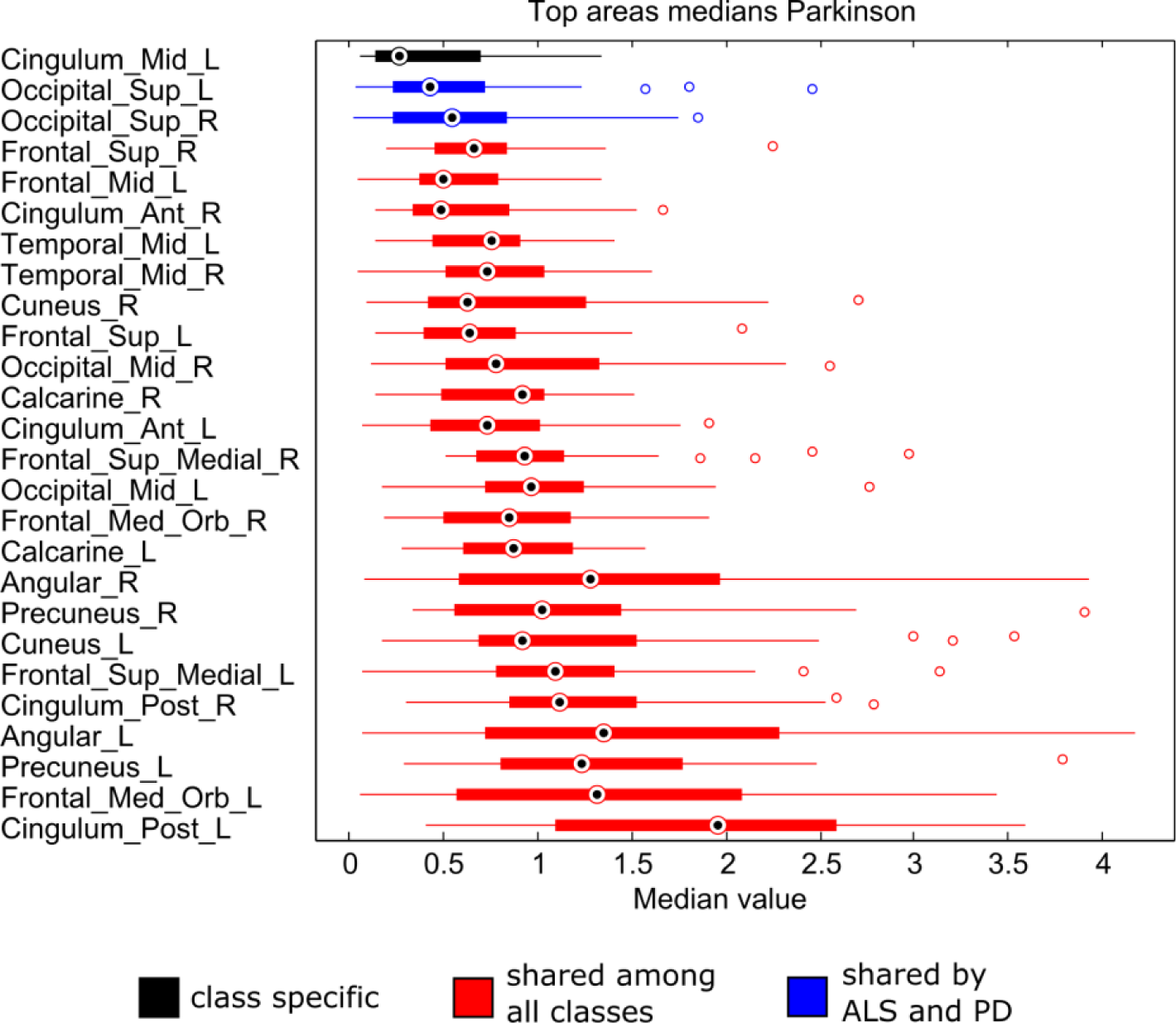
Box plots of the distribution of the medians of each of the selected anatomical areas across subjects of the PD class. The only class-specific region (in black) occupies the first rank, followed by the two regions shared with the ALS class (in blue).

**Figure 10.**
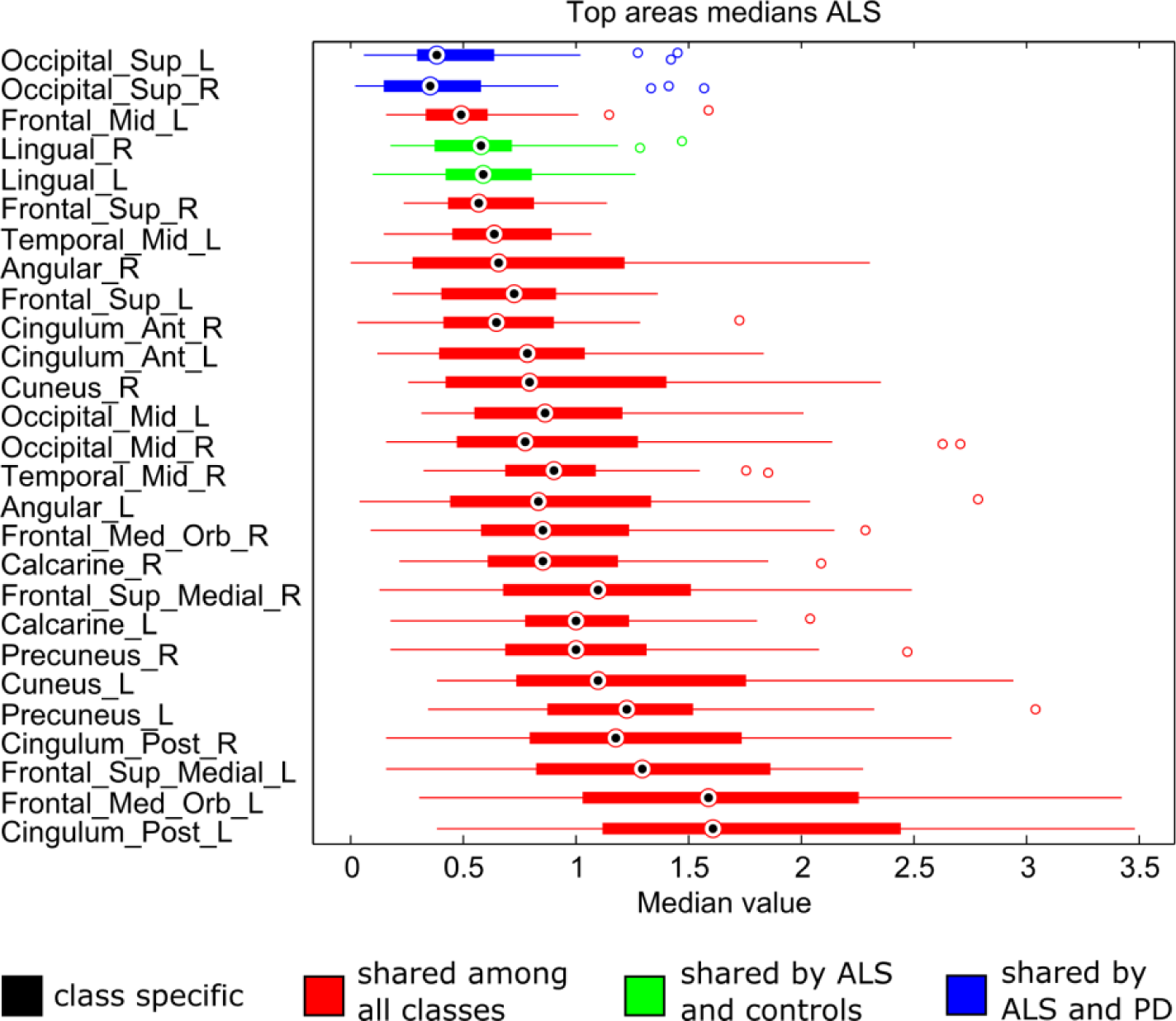
**1** Box plots of the distribution of the medians of each of the selected anatomical areas across subjects of the ALS class. There are no class-specific regions, but the ones shared with PD (in blue) and controls (in green) are listed in the top ranks with narrower distributions.

**Figure 11.**
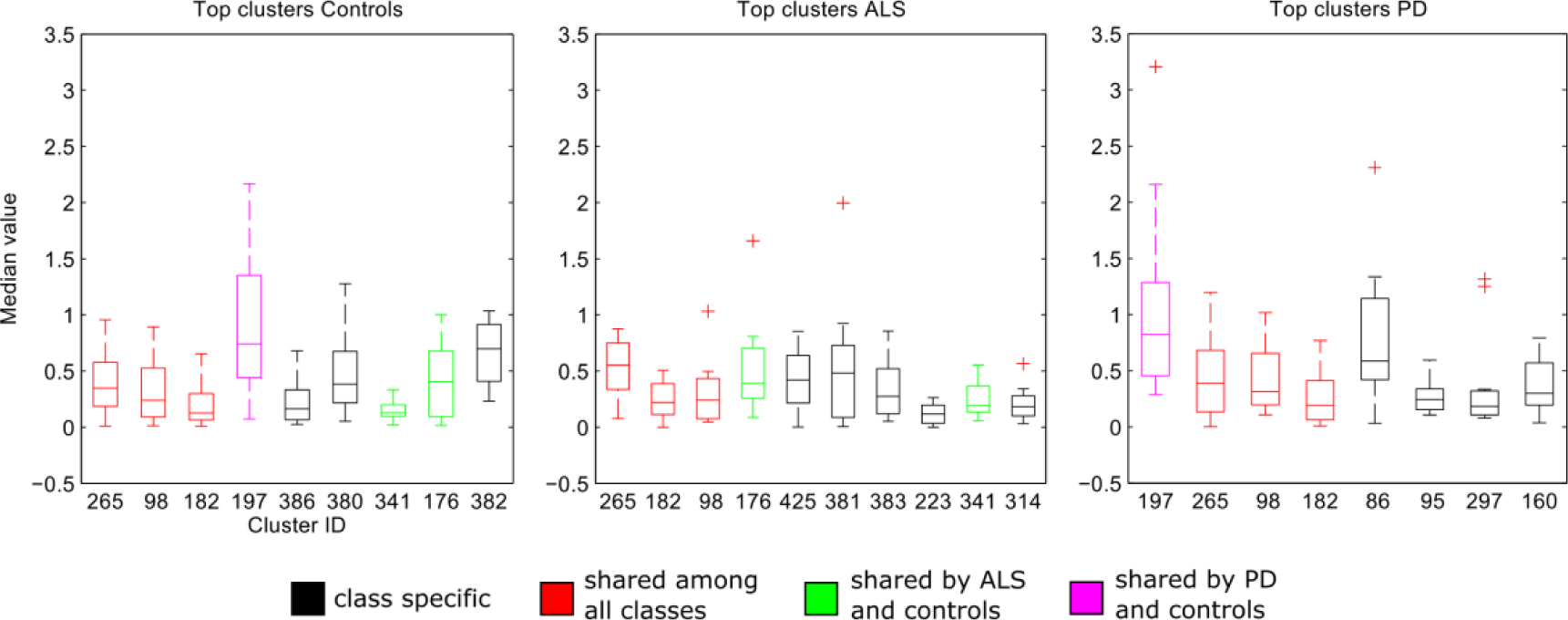
Box plots of the distribution of the medians of each of the selected clusters across subjects of the same class. In general, the parcels show a reduced variability compared to the anatomical regions, but this is not unexpected since the clusters are smaller and the selected parcels are fewer.

To assess the statistical significance of the results, we first computed the modified Kendall tau distance^3^ (Lin 2010) between the ranked lists of regions selected for each pair of conditions, to have a quantitative measure of how different the respective solutions are. We then estimated the empirical distribution of chance by repeating our analysis using 1000 random permutations of the phenotypes (i.e. we randomly assigned each subject to one of the three groups, keeping everything else the same) and computing the Kendall tau distances again for each of the permutations. Finally, we computed the p-value for each comparison by counting how many times the measured distance was greater in the random permutations than in the original analysis. Figure 12 shows the empirical distributions computed for each comparison together with the Kendall tau distance obtained in the original analysis. Multiple comparison correction was performed with the Benjamini-Hochberg procedure, separately for the results based on the anatomical parcellation and for those based on clustering. Significant differences (corrected p-value<0.05) were detected when comparing Controls vs. ALS and Controls vs. PD when using the anatomical parcellation, and ALS vs. PD when using clustering. Please note that the value of the Kendall distances computed between sets of anatomical areas and between clusters are not directly comparable, since they are defined in two different spaces (the 90 anatomical areas and the 405 clusters, respectively).

**Figure 12.**
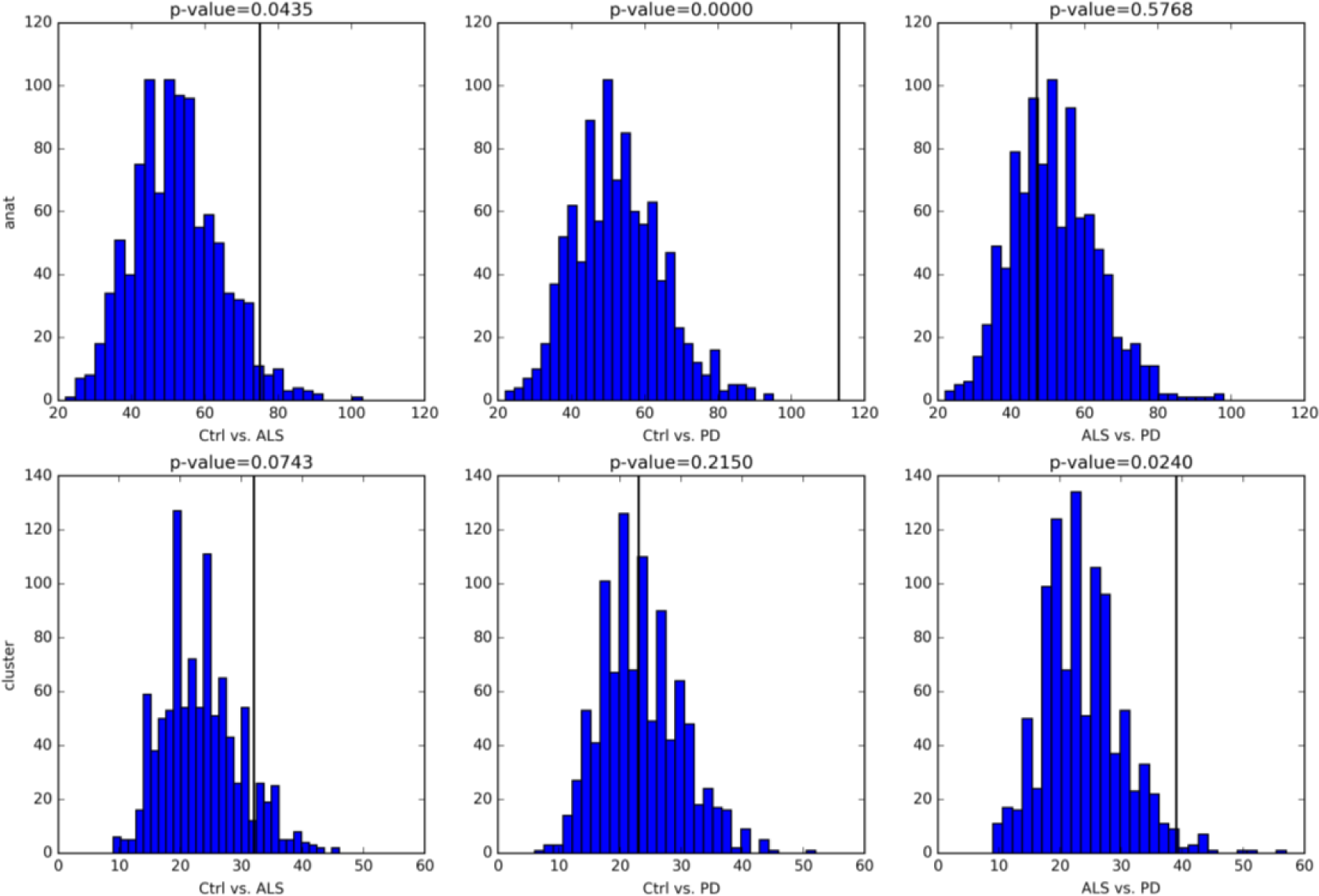
Empirical distributions of chance for the Kendall tau distance computed between ranked lists of regions selected for each pair of conditions (from left to right: Controls vs. ALS, Controls vs. PD, ALS vs. PD). Results obtained with the anatomical parcellation are showed in the top row, while results obtained with clustering are showed in the bottom row. The black vertical lines indicate the value of the Kendall tau distance computed in the original analysis. All comparisons are significant (p-value <0.05) except for ALS vs. PD when using the anatomical parcellation and Controls vs. PD when using clustering.

The stability of the results was assessed by repeating the same analysis on subgroups of subjects from each condition, specifically 1000 bootstrap samples were generated by sampling with replacement 2/3 of the initial groups of subjects. The similarity between two sets of regions selected for two subgroups from the same condition was measured with the Jaccard index, that is defined as the size of the intersection divided by the size of the union of the sets and is comprised between 0 (no overlap) and 1 (complete overlap). A final score per condition was computed as the average Jaccard index across all the comparisons. Since varying the initial set of subjects results in a different clustering for each subsample, when comparing sets of clusters the Jaccard index was computed between the sets of voxels constituting the clusters included in the solutions. Moreover, to verify if clusters were lying in the same anatomical regions, we identified the AAL parcels containing the clusters and computed the Jaccard index between sets of parcels. Results are reported in Table 3. A Wilcoxon rank-sum test indicated that all pair-wise differences were significant (p<0.001).

**Table 3.** For each condition, the average Jaccard index computed between pairs of solutions derived from 1000 subsamples of the original set of subjects is reported. The scores are higher for the anatomical areas than for clusters across conditions. The ALS solutions are the most stable across analyses.

| **Jaccard index** | **Controls** | **ALS** | **PD** |
| --- | --- | --- | --- |
| **AAL parcels** | $0.61 \pm 0.30$ | $0.91 \pm 0.04$ | $0.85 \pm 0.07$ |
| **clusters** | $0.19 \pm 0.06$ | $0.20 \pm 0.06$ | $0.15 \pm 0.07$ |
| **AAL parcels containing clusters** | $0.51 \pm 0.11$ | $0.58 \pm 0.10$ | $0.49 \pm 0.13$ |

## 4. Discussion

A novel framework based on clustering and stochastic rank aggregation has been evaluated using DMN maps from resting-state fMRI scans of ALS and PD patients and of healthy controls. As an alternative to clustering, a purely anatomical definition of brain parcels to extract regional DMN features for ranking was also considered. Since our goal was to demonstrate the utility of the proposed approach, we deemed out of the scope of this paper to extend the analysis to other resting state networks.

While in the clustering-based analyses about 2% of the clusters (8 or 9 out of 405) were selected in the final rankings, in the case of anatomical defined areas up to one third of the parcels (30 out of 90) are part of the solution. One reason for this might be that anatomical areas are bigger and fewer compared to clusters and since the median is used as representative feature this might flatten the differences across subjects resulting in more conforming rankings. This would also explain why more anatomical parcels are shared in the rankings across classes. Indeed, observing the Venn diagrams in figure 5, most of the anatomical areas are shared among the three classes, while there are more class-specific regions when using clusters. Another aspect to consider when interpreting the overlapping among selected regions is that for each subject the input data correspond to the independent component that best fits a template of the DMN. Therefore, it is reasonable to expect activations in approximately the same regions. It should also be noted that the number of overlapping regions is likely influenced by the size and number of input parcels. Indeed, working with larger (fewer) parcels might smooth out the underlying signal thus masking differences between classes and consequently resulting in more overlapping regions. However, the aim of aggregating rankings by class is precisely that of revealing patterns that are disease-specific: differences might be subtle and undetectable with a standard parametric test, but within-class coherence can provide further insights into disease-dependent alterations. Although not all between-group differences were significant, the results of permutation testing (figure 12) confirmed that the proposed approach can indeed highlight differences between conditions, thus providing a new method to search for a set of regional features characteristic of a specific neurological condition, i.e. what is usually called a neuromarker.

It is interesting to observe that the selected anatomical parcels cover most of the DMN, that is the spatial component that was extracted by ICA, meaning that although not very selective, the approach based on the anatomical parcellation led to the subset of regions that contribute to this network. As mentioned above, clusters are smaller than anatomical parcels and might therefore unveil differences at a finer granularity. Indeed, the example of figure 7, where a region included in the anatomical solutions of all three classes is compared to two clusters that are specific for the control group and one cluster that is specific for the ALS class, suggests that working with smaller regions might bring to light slight differences that are not evident at a higher level, because they are averaged out when considering a larger region. Furthermore, the fact that the clustering-based solution was able to detect a difference between the ALS and PD conditions, while the selected anatomical areas identified a contrast only when comparing patients to controls, could indicate that the data-driven approach is more sensitive to variations in the underlying signal that can differentiate pathological conditions. This could be explained considering that the clustering is built on functional data, while the anatomical parcellation is based on a priori information, hence specific aspects of the neurodegenerative conditions might be lost in this representation. Nevertheless, most of the clusters in the solution are included in one of the anatomical parcels (see last column of Table 2), often with a similar ranking between the two approaches, demonstrating the consistency of the results.

If we study in detail the clusters selected for each class, we can observe that clusters 86, 95 and 160, that are class specific for PD, all lie in Brodmann area 40, in the inferior parietal cortex, that has been shown to be relevant for this disease in previous fMRI studies: in Tessitore et al. (2012b) a decreased functional connectivity of the bilateral inferior parietal cortex in the DMN was observed in patients with PD; the results of Amboni et al. (2015) suggest that a functional disconnection of the frontoparietal network could be associated with mild cognitive impairment in PD. Another class specific cluster for PD is cluster 297 in Brodmann area 39 in the medial temporal lobe, whose relation with PD has been investigated in Gorges et al. (2013) and Tessitore et al. (2012b), where a decreased connectivity in the DMN was observed between this area and the posterior cingulate cortex, and with the prefrontal cortex, respectively.

Clusters 98 and 265, that are high-ranking in all three classes, lie in Brodmann area 10 in the prefrontal cortex, a region related to both ALS and PD: a weaker connectivity of the prefrontal region was observed in ALS patients in the DMN (Mohammadi et al. 2009; Agosta et al. 2013) and in the salience network (Trojsi et al. 2015); a deactivation of the medial prefrontal cortex was measured in PD patients in the DMN (Van Eimeren et al. 2009; Gorges et al. 2013) and in the fronto-parietal network (Amboni et al. 2015); Tessitore et al. (2012a) showed that PD patients with freezing of gait present reduced functional connectivity within the executive-attention network in the middle frontal gyrus. Considering the five clusters that are specific to the ALS class, we observe that clusters 381 and 383 are located in the temporal lobe; cluster 314 is included in Brodmann areas 18 (in the occipital lobe) and 7, in the left precuneus, that has been previously associated to ALS in the study by Agosta et al. (2013), where this region exhibited enhanced connectivity in the DMN; cluster 223 covers Brodmann areas 23 and 24, corresponding to the posterior and anterior cingulate cortex, mentioned in Tedeschi et al. (2012), where the DMN showed a disease-by-age interaction in the posterior cingulate cortex, and in Mohammadi et al. (2009) where the DMN showed less activation in ALS patients compared to controls; cluster 425 lies in Brodmann area 8 on the middle frontal gyrus, that has been associated with cognitive deficit in ALS patients in a PET study by Wicks et al. (2008), while in Terada et al. (2016) the gray matter volume measured within the right middle frontal gyrus in ALS patients was significantly lower than in healthy controls. If we observe the clusters that are class specific for control subjects, two of them (clusters 380 and 382) are on the middle temporal gyrus and one (cluster 386) on the fusiform gyrus. Both these regions are mentioned in studies on cortical thickness that investigated healthy aging as opposed to neurodegenerative disorders: in a work by Convit et al. (2000), the volumes of the fusiform gyrus and the middle (and inferior) temporal gyrus are shown to predict decline to Alzheimer’s disease (AD) in non-demented elderly; while in a work of Hänggi et al. (2011), the volume of the right middle temporal gyrus revealed promising diagnostic values to distinguish AD from mild cognitive impairment. Another study (Huettel et al. 2001) investigated the aging-related changes of the haemodynamic response in regions surrounding the fusiform gyrus. Finally, three clusters (182, 197 and 341) fall in regions of white matter and might therefore be resulting from noise.

From the point of view of the stability, the results based on the anatomical parcellation appear to be more stable (Table 3), however this is not unexpected for two reasons: first, the anatomical parcellation is the same across subsamples, while the clustering-based solutions, for their data-driven nature, change with every different subsample; second, as mentioned above, the selected anatomical areas tend to cover up to one third of the total number of parcels, while the selected clusters cover only about the 2% of the total number of clusters, hence a larger overlap is more likely with the anatomical areas. Additionally, considering the high-dimensional nature of the input data, the stability of the clustering-based results is also affected by the limited sample size; this issue could however be alleviated using larger data sets. Another aspect that might negatively affect stability is the choice of the clustering algorithm used to generate the initial parcellation; we adopted k-means but other methods might find more robust brain clusters (Thirion et al. 2014). Nevertheless, the bottom row of table 3 shows that although the selected voxels are not exactly the same across different subsamples of subjects, about a half of the identified anatomical regions are shared among solutions. Indeed, when evaluating clustering results, the most relevant aspect to consider is whether the information represented by clusters is consistently preserved, and the fact that the anatomical areas containing clusters in each subsample are overlapping is an indirect measure of how stable the information content of clusters is. It should be also pointed out that the essential role of the consensus clustering stage in the presented approach is to improve the selection of data entering the rank aggregation and the *TopKLists* algorithm, rather than provide a new (and stable) data-driven parcellation to be proposed as a generalizable alternative to the predefined anatomical parcellation. Indeed, as illustrated above, the use of clustering within the proposed pipeline increases the sensitivity of the rank aggregation method specifically in detecting (expectedly subtler) differences between two different pathological conditions, and such evidence remains valid even if the parcellation itself obtained via clustering is not very stable across possible subgroups and cannot be as stable as a pre-defined parcellation. In other words, the role of clustering in the presented approach (i.e. data selection for ranking) is somewhat similar to the role of principal component analysis (PCA) for subspace selection in group-level factor analyses. When performing factor analyses of high-dimensional data, the data are sometimes projected to a low-dimensional subspace (e.g. the first *k* principal components) and then the PCA scores are used for the factorial analyses. However, if repeating the same analysis on subgroups (e.g. to see whether the same group effects are significant or not) it is not necessarily relevant to see how similar the subspaces of different subgroups would be, especially if one would be interested to see how the group effects of interest (e.g. the difference between conditions) remain significant or not in each bootstrap sample.

Another interesting result of the stability analysis is that, regardless of the type of analysis, the ALS solutions are the most stable; this could suggest that the pattern of the activation within the DMN for this condition has a more distinctive signature and as a consequence there is greater consistency when comparing results between subsamples.

We also tried to validate our results using Neurosynth (Yarkoni et al. 2011). Note that these results are however limited to the terms/documents currently available in the Neurosynth database. For example, there are only 15 studies on ALS included in the text corpus, as opposed to the 104 available on PD. Moreover, the term ‘amyotrophic lateral sclerosis’ is not included in the list of features available for term-based meta-analysis. For this reason, we restricted part of the analyses to PD only. Since we used Talairach space and Neurosynth is based on the MNI space, we converted our coordinates using the same transformation used in the Neurosynth software. We performed two distinct analyses. The first was the reverse-inference analysis, to see if the coordinates of the selected regions per each class were associated to terms relevant to the studied conditions by previous literature (according to Pearson’s correlation between z-score activation maps, with FDR-adjusted p-value<0.05). This was done for all the terms available in the database, and for 200 topics derived from the terms applying Latent Dirichlet allocation (Poldrack et al. 2012). Table 4 and 5 report the results for terms and topics, respectively.

**Table 4.**
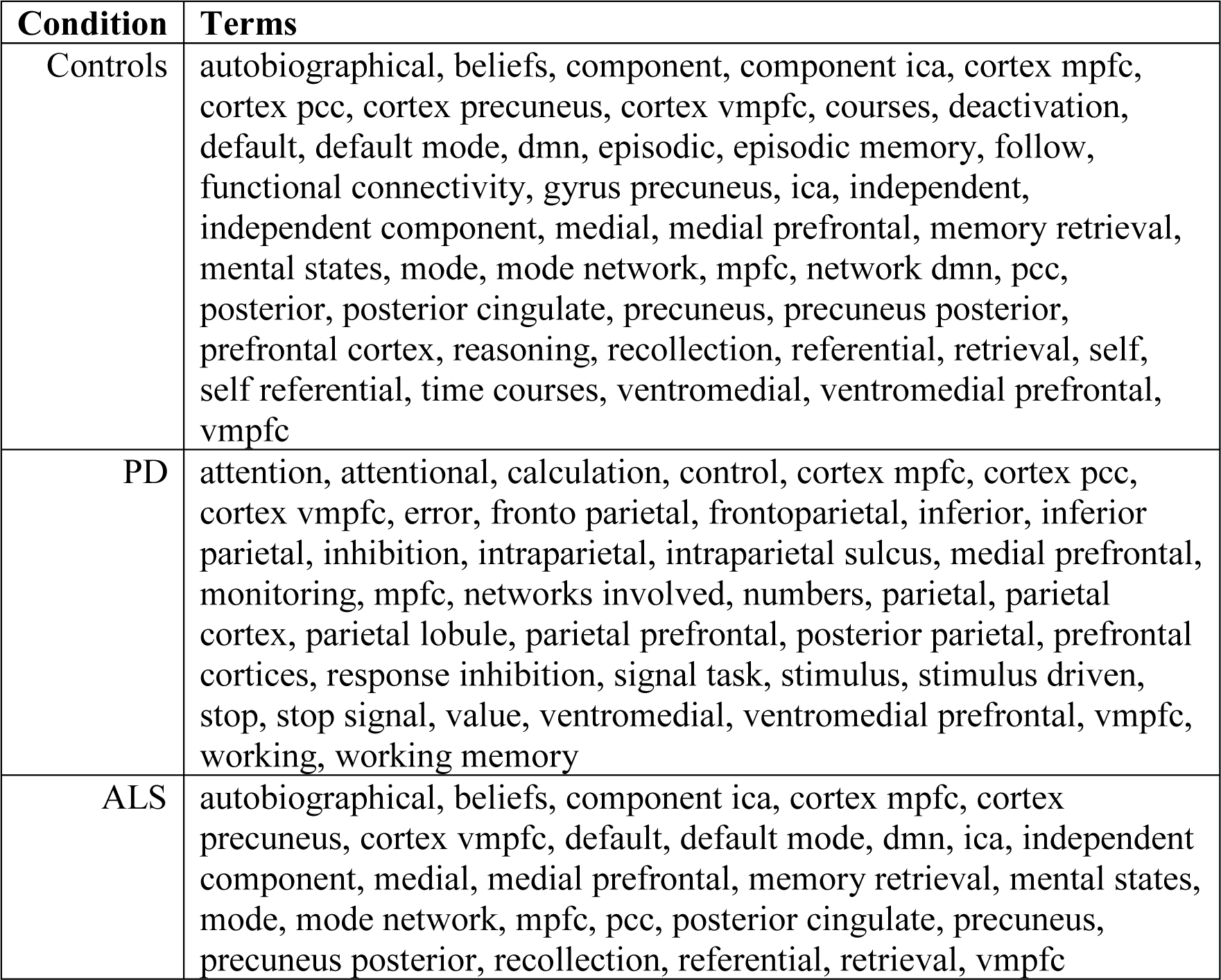
List of terms resulting from the reverse-inference analysis for each condition. Each of the term listed for each class was used in studies that reported an activation in the regions selected for that condition.

**Table 5.**
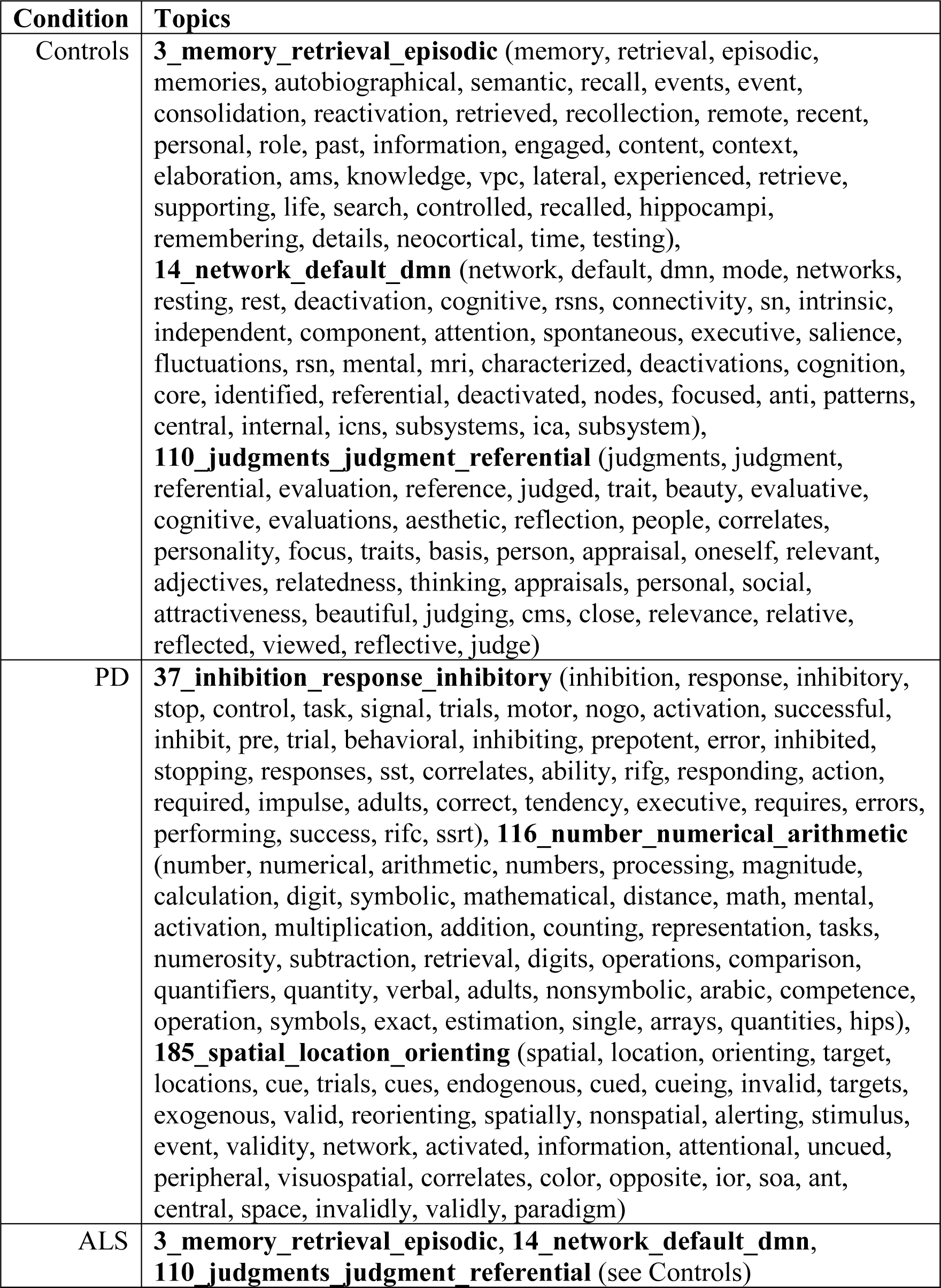
List of topics associated with the selected regions for each class. The terms defining each topic are listed in brackets.

From table 4 we can observe that the retrieved terms are more generic than disease-specific. They mostly refer to brain regions (e.g., precuneus, posterior cingulate, parietal cortex), ICA (e.g., component, independent componenct, ica), the default mode network (e.g., default, default mode, dmn), and functions associated with the default mode network (autobiographical, episodic memory, recollection, memory retrieval, beliefs) (Buckner et al. 2008). This replicates also in the topic-based analysis (table 5) for the controls and ALS classes, where the retrieved topic can all be associated to the DMN and its function (3_memory_retrieval_episodic, 14_network_default_dmn, 10_judgments_judgment_referential). However, for the PD class, there are some additional terms and topics not shared with the other two classes, referring to executive functions (attention, calculation, control, working memory, numbers, 116_number_numerical_arithmetic), inhibition (37_inhibition_response_inhibitory), and visuospatial orienting (185_spatial_location_orienting). All these three domains have been previously linked to PD (Monchi et al. 2007; Beato et al. 2008; Possin et al. 2008; Pereira et al. 2009; McKinlay et al. 2010; Disbrow et al. 2013; Dirnberger and Jahanshahi 2013; Jahanshahi et al. 2015; Di Rosa et al. 2017).

A second analysis consisted in computing a map of activations derived from the meta-analysis of studies associated with the term “Parkinson” (this was not possible for ALS). We then looked at the overlapping between the regions selected with our approach (1315 activations) and the 4135 activations resulting from the meta-analysis. Although the voxels in the intersection were only 19, all activations were found in Brodmann area 40 (inferior parietal cortex), that is the region in which three out of the four clusters specific to the PD class were found, and that was reported to be overactive in PD patients performing motor tasks (Samuel et al. 1997; Hanakawa et al. 1999).

## 5. Conclusion

We presented a novel methodology to automatically detect regions that show a common behaviour in a class of subjects. Looking for commonalities instead of differences between groups is advantageous because the actual differences might be masked by noise of different origins. This approach takes into account both inter-subject variability and noise by excluding from the analysis brain parcels whose patterns of activation are incoherent across subjects of the same diagnostic group, while retaining regions for which a sufficient degree of consensus exists. The further advantage of combining this technique with a clustering-based parcellation is that of making the whole approach data-driven and hence more sensitive to subtle variations in the data that are however consistent across subjects of the same class. In this work, the proposed methodology was applied on DMN maps derived from resting-state fMRI acquisitions of healthy controls, PD and ALS patients, and results are consistent with previous literature, thus indicating that this approach might be a suitable method to search for a set of regional features characteristic of a specific neurological condition, i.e. what is usually called a neuromarker, which is a fundamental step in the development of fully automated data-driven techniques to support early diagnoses of neurodegenerative diseases.

Nonetheless, the same framework can be applied with parcels derived from pre-existing brain atlases. As a proof of concept, we showed an example of application of our method in combination with the AAL atlas, because it is the most commonly adopted in literature and our goal was to facilitate the comparison of findings. Our results suggest that subtler differences can be uncovered when using smaller, more localized regions of interest: compared to the results obtained with the anatomical parcellation, the regions identified through clustering were able to highlight differences between neurological conditions better than between healthy controls and patients. However, these results are not conclusive, and further experiments comparing a range of healthy and neurological conditions (as well as statistal tests on new data) would be needed to confirm that this holds in general. In addition, to prove the superiority of a data-driven parcellation over a predefined atlas it would be necessary to replicate this analysis on other datasets and to extend the study to other atlantes. A promising candidate would be the multi-modal brain parcellation recently proposed by Glasser et al. (2016), yet this approach requires different imaging modalities that were not available for the dataset under study.

Another limitation of this study is that we did not explicitly explore the effect of the resolution of the brain parcellation on the stability of the final rankings, although we partly addressed the effect of the choice of the parcellation on the outcome of the analysis. While it is likely that working with larger regions (fewer parcels) would increase the stability of the final rankings as in the case of the anatomical parcellation, this is not necessarily an advantage since averaging the signal over a large parcel might flatten some of the differences that might be relevant to characterize different neurological conditions. A systematic study on the effect of the number of parcels in fMRI data analyses has previously shown that the number of clusters (k) can greatly affect the accuracy and reproducibility of the results and that k=200 is a lower bound to optimally represent the variability of functional information in fMRI data (Thirion et al. 2014).

In future work, it would be interesting to apply the same method to task fMRI data, where it is easier to detect activation loci and more prior knowledge is available to define regions of interest. This would allow observing how the rankings of regions vary between healthy subjects and patients in the performance of specific tasks.

## Conflict of Interest

The authors declare that they have no conflict of interest.

## Information Sharing Statement

Source code (Matlab and R scripts) implementing the methods described in this paper, as well as the preprocessed (and anonymized) group data sets used in the analyses, are available on GitHub (https://github.com/paola-g/STRAIN).

## Footnotes

1 The template binary mask was computed as the mean DMN map of a separate population of control subjects.

2 A second assumption is that *p_i_* decreases when *i* increases.

3 The original version of the Kendall tau distance simply counts the number of pairwise discordances between the two ranked lists defined on the same sets of objects; the modified version can deal also with partial lists.

## References

Agosta F, Canu E, Valsasina P, et al (2013) Divergent brain network connectivity in amyotrophic lateral sclerosis. Neurobiol. Aging 34:419–27. doi: 10.1016/j.neurobiolaging.2012.04.015

Amboni M, Tessitore A, Esposito F, et al (2015) Resting-state functional connectivity associated with mild cognitive impairment in Parkinson’s disease. J. Neurol. 262:425–434. doi: 10.1007/s00415-014-7591-5

Amelio A, Pizzuti C (2015) Is Normalized Mutual Information a Fair Measure for Comparing Community Detection Methods? In: Proceedings of the 2015 IEEE/ACM International Conference on Advances in Social Networks Analysis and Mining 2015 - ASONAM ‘15. ACM Press, New York, New York, USA, pp 1584–1585

Beato R, Levy R, Pillon B, et al (2008) Working memory in Parkinson’s disease patients: clinical features and response to levodopa. Arq. Neuropsiquiatr. 66:147–51.

Bosch OG, Esposito F, Dornbierer D, et al (2018) Gamma-hydroxybutyrate increases brain resting-state functional connectivity of the salience network and dorsal nexus in humans. Neuroimage 173:448–459. doi: 10.1016/j.neuroimage.2018.03.011

Buckner RL, Andrews-Hanna JR, Schacter DL (2008) The Brain’s Default Network. Ann. N. Y. Acad. Sci. 1124:1–38. doi: 10.1196/annals.1440.011

Chiò A, Pagani M, Agosta F, et al (2014) Neuroimaging in amyotrophic lateral sclerosis: insights into structural and functional changes. Lancet Neurol. 13:1228–1240. doi: 10.1016/S1474-4422(14)70167-X

Convit A, De Asis J, De Leon MJ, et al (2000) Atrophy of the medial occipitotemporal, inferior, and middle temporal gyri in non-demented elderly predict decline to Alzheimer’s disease. Neurobiol. Aging 21:19–26. doi: 10.1016/S0197-4580(99)00107-4

Damoiseaux JS, Rombouts SARB, Barkhof F, et al (2006) Consistent resting-state networks across healthy subjects. Proc. Natl. Acad. Sci. U. S. A. 103:13848–53. doi: 10.1073/pnas.0601417103

De Micco R, Tessitore A, Paccone A, et al (2013) Dopaminergic modulation of the resting-state sensori-motor network in drug-naive patients with Parkinson’s disease. Mov. Disord. 28:S66–S66.

Di Rosa E, Pischedda D, Cherubini P, et al (2017) Working memory in healthy aging and in Parkinson’s disease: evidence of interference effects. Aging, Neuropsychol. Cogn. 24:281–298. doi: 10.1080/13825585.2016.1202188

Dirnberger G, Jahanshahi M (2013) Executive dysfunction in Parkinson’s disease: A review. J. Neuropsychol. 7:193–224. doi: 10.1111/jnp.12028

Disbrow EA, Sigvardt KA, Franz EA, et al (2013) Movement activation and inhibition in Parkinson’s disease: a functional imaging study. J. Parkinsons. Dis. 3:181–92. doi: 10.3233/JPD-130181

Eklund A, Nichols TE, Knutsson H (2016) Cluster failure: Why fMRI inferences for spatial extent have inflated false-positive rates. Proc Natl Acad Sci. doi: 10.1073/pnas.1602413113

Esposito F, Aragri A, Pesaresi I, et al (2008) Independent component model of the default-mode brain function: combining individual-level and population-level analyses in resting-state fMRI. Magn. Reson. Imaging 26:905–913. doi: 10.1016/J.MRI.2008.01.045

Esposito F, Goebel R (2011) Extracting functional networks with spatial independent component analysis: the role of dimensionality, reliability and aggregation scheme. Curr. Opin. Neurol. 24:378–85. doi: 10.1097/WCO.0b013e32834897a5

Esposito F, Pignataro G, Di Renzo G, et al (2010) Alcohol increases spontaneous BOLD signal fluctuations in the visual network. Neuroimage 53:534–543. doi: 10.1016/j.neuroimage.2010.06.061

Esposito F, Scarabino T, Hyvarinen A, et al (2005) Independent component analysis of fMRI group studies by self-organizing clustering. Neuroimage 25:193–205. doi: 10.1016/j.neuroimage.2004.10.042

Esposito F, Tessitore A, Giordano A, et al (2013) Rhythm-specific modulation of the sensorimotor network in drug-naïve patients with Parkinson’s disease by levodopa. Brain 136:710–725. doi: 10.1093/brain/awt007

Galdi P, Fratello M, Trojsi F, et al (2017) Consensus-based feature extraction in rs-fMRI data analysis. Soft Comput. 1–11. doi: 10.1007/s00500-017-2596-5

Gattellaro G, Minati L, Grisoli M, et al (2009) White matter involvement in idiopathic Parkinson disease: a diffusion tensor imaging study. AJNR. Am. J. Neuroradiol. 30:1222–6. doi: 10.3174/ajnr.A1556

Glasser MF, Coalson TS, Robinson EC, et al (2016) A multi-modal parcellation of human cerebral cortex. Nature 536:171–178. doi: 10.1038/nature18933

Gorges M, Müller H-P, Lulé D, et al (2013) Functional connectivity within the default mode network is associated with saccadic accuracy in Parkinson’s disease: a resting-state FMRI and videooculographic study. Brain Connect. 3:265–72. doi: 10.1089/brain.2013.0146

Goutte C (1999) On clustering fMRI time series.

Goutte C, Hansen LK, Liptrot MG, Rostrup E (2001) Feature-space clustering for fMRI meta-analysis. Hum. Brain Mapp. 13:165–183. doi: 10.1002/hbm.1031

Greicius MD, Flores BH, Menon V, et al (2007) Resting-State Functional Connectivity in Major Depression: Abnormally Increased Contributions from Subgenual Cingulate Cortex and Thalamus. Biol. Psychiatry 62:429–437. doi: 10.1016/j.biopsych.2006.09.020

Greicius MD, Krasnow B, Reiss AL, Menon V (2003) Functional connectivity in the resting brain: a network analysis of the default mode hypothesis. PNAS 100:253–8. doi: 10.1073/pnas.0135058100

Greicius MD, Srivastava G, Reiss AL, Menon V (2004) Default-mode network activity distinguishes Alzheimer’s disease from healthy aging: evidence from functional MRI. Proc. Natl. Acad. Sci. U. S. A. 101:4637–42. doi: 10.1073/pnas.0308627101

Hall P, Schimek MG (2012) Moderate-Deviation-Based Inference for Random Degeneration in Paired Rank Lists. J. Am. Stat. Assoc. 107:661–672. doi: 10.1080/01621459.2012.682539

Hanakawa T, Fukuyama H, Katsumi Y, et al (1999) Enhanced lateral premotor activity during paradoxical gait in parkinson’s disease. Ann. Neurol. 45:329–336. doi: 10.1002/1531-8249(199903)45:3<329::AID-ANA8>3.0.CO;2-S

Hänggi J, Streffer J, Jäncke L, Hock C (2011) Volumes of lateral temporal and parietal structures distinguish between healthy aging, mild cognitive impairment, and Alzheimer’s disease. J. Alzheimers. Dis. 26:719–34. doi: 10.3233/JAD-2011-101260

Harrison BJ, Pujol J, Lopez-Sola M, et al (2008) Consistency and functional specialization in the default mode brain network. Proc. Natl. Acad. Sci. 105:9781–9786. doi: 10.1073/pnas.0711791105

Hepp DH, Foncke EMJ, Olde Dubbelink KTE, et al (2017) Loss of Functional Connectivity in Patients with Parkinson Disease and Visual Hallucinations. Radiology 285:896–903. doi: 10.1148/radiol.2017170438

Herrington TM, Briscoe J, Eskandar E (2017) Structural and Functional Network Dysfunction in Parkinson Disease. Radiology 285:725–727. doi: 10.1148/radiol.247172401

Heuvel M van den, Mandl R, Pol HH (2008) Normalized Cut Group Clustering of Resting-State fMRI Data. PLoS One 3:e2001. doi: 10.1371/JOURNAL.PONE.0002001

Huettel SA, Singerman JD, McCarthy G (2001) The Effects of Aging upon the Hemodynamic Response Measured by Functional MRI. Neuroimage 13:161–175. doi: 10.1006/nimg.2000.0675

Hyvarinen A (1999) Fast and Robust Fixed-Point Algorithm for Independent Component Analysis. IEEE Trans. Neur. Net. 10:626–634.

Hyvärinen A, Oja E (2000) Independent component analysis: algorithms and applications. Neural networks 13:411–430.

Iyer PM, Egan C, Pinto-Grau M, et al (2015) Functional connectivity changes in resting-state EEG as potential biomarker for Amyotrophic Lateral Sclerosis. PLoS One. doi: 10.1371/journal.pone.0128682

Jahanshahi M, Obeso I, Baunez C, et al (2015) Parkinson’s Disease, the Subthalamic Nucleus, Inhibition, and Impulsivity. Mov. Disord. 30:128–140. doi: 10.1002/mds.26049

Kiernan MC, Vucic S, Cheah BC, et al (2011) Amyotrophic lateral sclerosis. Lancet 377:942–955. doi: 10.1016/S0140-6736(10)61156-7

Lin S (2010) Rank aggregation methods. Wiley Interdiscip. Rev. Comput. Stat. 2:555–570. doi: 10.1002/wics.111

Lin S, Ding J (2009) Integration of Ranked Lists via Cross Entropy Monte Carlo with Applications to mRNA and microRNA Studies on JSTOR. In: Biometrics. http://www.jstor.org/stable/25502239?seq=1#page_scan_tab_contents. Accessed 7 Jan 2016

Lomen-Hoerth C, Murphy J, Langmore S, et al (2003) Are amyotrophic lateral sclerosis patients cognitively normal? Neurology 60:1094–1097. doi: 10.1212/01.WNL.0000055861.95202.8D

Luo C, Chen Q, Huang R, et al (2012) Patterns of Spontaneous Brain Activity in Amyotrophic Lateral Sclerosis: A Resting-State fMRI Study. PLoS One 7:e45470. doi: 10.1371/journal.pone.0045470

Luo C, Guo X, Song W, et al (2015) The trajectory of disturbed resting-state cerebral function in Parkinson’s disease at different Hoehn and Yahr stages. Hum. Brain Mapp. 36:3104–3116. doi: 10.1002/hbm.22831

McKinlay A, Grace RC, Dalrymple-Alford JC, Roger D (2010) Characteristics of executive function impairment in Parkinsons disease patients without dementia. J. Int. Neuropsychol. Soc. 16:268–277. doi: 10.1017/S1355617709991299

Meilă M (2007) Comparing clusterings—an information based distance. J. Multivar. Anal. 98:873–895. doi: 10.1016/J.JMVA.2006.11.013

Menke RAL, Agosta F, Grosskreutz J, et al (2017) Neuroimaging Endpoints in Amyotrophic Lateral Sclerosis. Neurotherapeutics 14:11–23. doi: 10.1007/s13311-016-0484-9

Mohammadi B, Kollewe K, Samii A, et al (2009) Changes of resting state brain networks in amyotrophic lateral sclerosis. Exp. Neurol. 217:147–153. doi: 10.1016/j.expneurol.2009.01.025

Monchi O, Petrides M, Mejia-Constain B, Strafella AP (2007) Cortical activity in Parkinson’s disease during executive processing depends on striatal involvement. Brain 130:233–244. doi: 10.1093/brain/awl326

Olde Dubbelink KTEE, Hillebrand A, Stoffers D, et al (2014) Disrupted brain network topology in Parkinson’s disease: a longitudinal magnetoencephalography study. Brain 137:197–207. doi: 10.1093/brain/awt316

Pereira JB, Junqué C, Martí MJ, et al (2009) Neuroanatomical substrate of visuospatial and visuoperceptual impairment in Parkinson’s disease. Mov. Disord. 24:1193–1199. doi: 10.1002/mds.22560

Poldrack RA, Mumford JA, Schonberg T, et al (2012) Discovering Relations Between Mind, Brain, and Mental Disorders Using Topic Mapping. PLoS Comput. Biol. 8:e1002707. doi: 10.1371/journal.pcbi.1002707

Possin KL, Filoteo JV, Song DD, Salmon DP (2008) Spatial and object working memory deficits in Parkinson’s disease are due to impairment in different underlying processes. Neuropsychology 22:585–95. doi: 10.1037/a0012613

Raichle ME (2015) The Brain’s Default Mode Network. Annu. Rev. Neurosci. 38:433–447. doi: 10.1146/annurev-neuro-071013-014030

Samuel M, Ceballos-Baumann AO, Blin J, et al (1997) Evidence for lateral premotor and parietal overactivity in Parkinson’s disease during sequential and bimanual movements. A PET study. Brain 120:963–976. doi: 10.1093/brain/120.6.963

Schimek MG, Budinská E, Kugler KG, et al (2015) TopKLists: a comprehensive R package for statistical inference, stochastic aggregation, and visualization of multiple omics ranked lists. Stat. Appl. Genet. Mol. Biol. 14:311–6. doi: 10.1515/sagmb-2014-0093

Schimek MG, Myšičková A, Budinská E (2012) An Inference and Integration Approach for the Consolidation of Ranked Lists. Commun. Stat.-Simul. Comput. 41:1152–1166. doi: 10.1080/03610918.2012.625843

Smith SM, Fox PT, Miller KL, et al (2009) Correspondence of the brain’s functional architecture during activation and rest. Proc. Natl. Acad. Sci. 106:13040–13045. doi: 10.1073/pnas.0905267106

Suo X, Lei D, Li N, et al (2017) Functional Brain Connectome and Its Relation to Hoehn and Yahr Stage in Parkinson Disease. Radiology 285:904–913. doi: 10.1148/radiol.2017162929

Tedeschi G, Esposito F (2009) Neuronal Networks Observed with Resting State Functional Magnetic Resonance Imaging in Clinical Populations. Neuroimaging - Cogn Clin Neurosci. doi: 10.5772/23290

Tedeschi G, Trojsi F, Tessitore A, et al (2012) Interaction between aging and neurodegeneration in amyotrophic lateral sclerosis. Neurobiol. Aging 33:886–898. doi: 10.1016/j.neurobiolaging.2010.07.011

Terada T, Obi T, Miyata J, et al (2016) Correlation of frontal atrophy with behavioral changes in amyotrophic lateral sclerosis. Neurol. Clin. Neurosci. 4:85–92. doi: 10.1111/ncn3.12046

Tessitore A, Amboni M, Esposito F, et al (2012a) Resting-state brain connectivity in patients with Parkinson’s disease and freezing of gait. Parkinsonism Relat. Disord. 18:781–787. doi: 10.1016/j.parkreldis.2012.03.018

Tessitore A, Esposito F, Vitale C, et al (2012b) Default-mode network connectivity in cognitively unimpaired patients with Parkinson disease. Neurology 79:2226–32. doi: 10.1212/WNL.0b013e31827689d6

Tessitore A, Giordano A, De Micco R, et al (2014) Sensorimotor Connectivity in Parkinsonâ€TMs Disease: The Role of Functional Neuroimaging. Front. Neurol. 5:180. doi: 10.3389/fneur.2014.00180

Thirion B, Flandin G, Pinel P, et al (2006) Dealing with the shortcomings of spatial normalization: Multi-subject parcellation of fMRI datasets. Hum. Brain Mapp. 27:678–693. doi: 10.1002/hbm.20210

Trojsi F, Esposito F, de Stefano M, et al (2015) Functional overlap and divergence between ALS and bvFTD. Neurobiol. Aging 36:413–423. doi: 10.1016/j.neurobiolaging.2014.06.025

Tzourio-Mazoyer N, Landeau B, Papathanassiou D, et al (2002) Automated anatomical labeling of activations in SPM using a macroscopic anatomical parcellation of the MNI MRI single-subject brain.

Van Eimeren T, Monchi O, Ballanger B, Strafella AP (2009) Dysfunction of the default mode network in Parkinson disease: a functional magnetic resonance imaging study. Arch. Neurol. 66:877–83. doi: 10.1001/archneurol.2009.97

Verstraete E, Foerster BR (2015) Neuroimaging as a New Diagnostic Modality in Amyotrophic Lateral Sclerosis. Neurotherapeutics 12:403–416. doi: 10.1007/s13311-015-0347-9

Verstraete E, Veldink JH, van den Berg LH, van den Heuvel MP (2014) Structural brain network imaging shows expanding disconnection of the motor system in amyotrophic lateral sclerosis. Hum. Brain Mapp. 35:1351–1361. doi: 10.1002/hbm.22258

Vinh NX, Epps J (2009) A Novel Approach for Automatic Number of Clusters Detection in Microarray Data Based on Consensus Clustering. In: 2009 Ninth IEEE International Conference on Bioinformatics and BioEngineering. IEEE, pp 84–91

Vinh NX, Epps J, Bailey J (2009) Information theoretic measures for clusterings comparison. In: Proceedings of the 26th Annual International Conference on Machine Learning - ICML ’09. pp 1–8

von Luxburg U (2010) Clustering Stability: An Overview. Found. Trends^®^ Mach. Learn. 2:235–274. doi: 10.1561/2200000008

Whitfield-Gabrieli S, Ford JM (2012) Default Mode Network Activity and Connectivity in Psychopathology. Annu. Rev. Clin. Psychol. 8:49–76. doi: 10.1146/annurev-clinpsy-032511-143049

Wicks P, Turner MR, Abrahams S, et al (2008) Neuronal loss associated with cognitive performance in amyotrophic lateral sclerosis: An (11 C)-flumazenil PET study. Amyotroph. Lateral Scler. 9:43–49. doi: 10.1080/17482960701737716

Yarkoni T, Poldrack RA, Nichols TE, et al (2011) Large-scale automated synthesis of human functional neuroimaging data. Nat. Methods 8:665–670. doi: 10.1038/nmeth.1635

